# Microgravity Accelerates Skeletal Muscle Degeneration: Functional and Transcriptomic Insights from a Muscle Lab-on-Chip Model Onboard the ISS

**DOI:** 10.1101/2025.01.26.634580

**Authors:** Maddalena Parafati, Zon Thwin, Legrand K. Malany, Paul M. Coen, Siobhan Malany

## Abstract

Microgravity accelerates skeletal muscle degeneration, mimicking aging, yet its effects on human muscle cell function and signaling remain underexplored. Using a muscle lab-on-chip model onboard the International Space Station, we examined how microgravity and electrically stimulated contractions influence muscle biology and age-related muscle changes. Our 3D bioengineered muscle model, cultured for 21 days (12 days in microgravity), included myobundles from young, active and older, sedentary individuals, with and without electrically stimulated contraction. Real-time data collected within an autonomous Space Tango CubeLab^TM^ showed reduced contraction magnitude in microgravity. Global transcriptomic analysis revealed increased gene expression and particularly mitochondrial-related gene expression in microgravity for the electrically stimulated younger myobundles, while the older myobundles were less responsive. Moreover, a comparative analysis using a skeletal muscle aging gene expression database revealed that certain age-induced genes showed changes in expression in myobundles from the younger cohort when exposed to microgravity, whereas these genes remained unchanged in myobundles from the older cohort. Younger, electrically stimulated myobundles in microgravity exhibited higher expression of 45 aging genes involved in key aging pathways related to inflammation and immune function, mitochondrial dysfunction, and cellular stress; and decreased expression of 41 aging genes associated with inflammation, and cell growth. This study highlights a unique age-related molecular signature in muscle cells exposed to microgravity and underscores electrical stimulation as a potential countermeasure. These insights advance understanding of skeletal muscle aging and microgravity-induced degeneration, informing strategies for mitigating age-related muscle atrophy in space and on Earth.

## INTRODUCTION

With the rise of human space exploration, research has focused on understanding how space travel affects human health, the progression of diseases, and the body’s mechanisms for maintaining homeostasis. Extended spaceflight missions cause various health issues, with multisystemic dysfunction being a primary concern (Krittanawong et al., 2022; Strollo, 1999; Vernikos and Schneider, 2010). The musculoskeletal system is particularly vulnerable to the effects of spaceflight. Prolonged muscle disuse and the absence of gravity-induced mechanical loading result in rapid muscle atrophy (Comfort et al., 2021; Juhl et al., 2021; Lee et al., 2022; Trappe et al., 2009; Vandenburgh et al., 1999; Zhang et al., 2018). It is well established that skeletal muscle adapts to changes in its mechanical environment by altering gene expression (Leuchtmann et al., 2021; Powers and Schrager, 2022; Tidball, 2005). Accelerated muscle atrophy occurring during spaceflight has been proposed to be a relevant model for understanding sarcopenia, the progressive muscle loss primarily driven by aging (Cannavo et al., 2022; Larsson et al., 2019). Astronauts exposed to microgravity experience up to a 30% reduction in skeletal muscle mass and strength within just one month of spaceflight (Williams et al., 2009). This decline not only hinders astronauts’ ability to perform essential tasks aboard the International Space Station (ISS) but also increases their risk of injury upon returning to Earth’s gravity. Without daily, intensive exercise, muscle mass in astronauts spending six months in microgravity would resemble that of an 80-year-old (Trappe et al., 2009). Investigating the mechanisms behind spaceflight-induced muscle degeneration offers valuable insights into processes underlying age-related muscle degeneration on Earth.

Microgravity affects biology not only at the tissue level in vivo, leading to muscle atrophy, but in a cell autonomous manner. The growing interest in studying the effects of microgravity on cells has spurned the development of innovative lab-on-chip technologies to facilitate these investigations. Previously, we demonstrated the utility of our 3D skeletal muscle lab-on-chip model for studying muscle biology and stress responses in an autonomous payload CubeLab^TM^ custom designed for spaceflight (Parafati et al., 2023). In the CubeLab^TM^, donor-derived myoblasts undergo myogenesis, fusing and responding to chemical cues to form functional, mature human myobundles. Studies have shown that electrical stimulation, which simulates muscle contraction during exercise, enhances the alignment and responsiveness of muscle fibers (Khodabukus et al., 2019; Nikolić et al., 2012; Zhao et al., 2020). Building on our previous mission to the ISS, we cultured myotubes with well-organized sarcomeres and contractile responses to electrical pulse sequences in the CubeLab^TM^ environment, validating functional response with previous studies (Giza et al., 2022).

The current study investigated how microgravity influences myocyte contractile function and gene expression, and the protective effects of electrical stimulation in 3D-engineered myobundles derived from muscle biopsies of younger and older participants. We sought to identify new age-related transcriptional signatures associated with muscle cell exposure to microgravity. We compared donor-derived gene expression signatures by analyzing the transcriptional changes in myobundles exposed to microgravity, with or without electrical stimulation. We hypothesize that the younger donor-derived myobundles would exhibit more pronounced transcriptional adaptations to microgravity compared to myobundles derived from older donors (Giza et al., 2022) and that electrical stimulation would partially mitigate these effects. Gene enrichment analysis revealed key signaling networks and regulatory mechanisms associated with microgravity exposure in myobundles. These findings advance our understanding of muscle health in both space and aging contexts and pave the way for developing targeted countermeasures against muscle atrophy caused by spaceflight and sarcopenia on Earth.

## RESULTS

### Skeletal muscle tissue chip payload integration for spaceflight

Skeletal muscle myobundles were derived from precursor cells isolated from muscle biopsies of young active (YA) and old sedentary (OS) participants, as previously described(Giza et al., 2022). The cells were embedded in a Matrigel-collagen hydrogel mixture and cultured within custom-designed microfluidic chips compatible with the spaceflight payload hardware(Parafati et al., 2023). An advancement in this mission compared to our previous ISS study was the incorporation of electrical stimulation (E-Stim) for a subset of YA and OS-derived myobundles to simulate muscle contraction over one week. The remaining myobundles were not electrically stimulated (No E-Stim), enabling a comparative analysis of global gene expression changes across spaceflight and ground control conditions as well as between E-Stim and No E-Stim treatments.

To facilitate electrical stimulation, we modified the tissue chip design by fabricating grooves in the polydimethylsiloxane (PDMS) structure to secure platinum electrodes along the outer edge of the media channel, positioned parallel to the aligned myobundles (Figure 1A). The chips were integrated into a custom fluid handling manifold. This manifold housed eight tissue chips with electrodes (four YA and four OS) on one side, connected to a pulse board, and eight chips without electrodes (four YA and four OS) on the opposite side (Figure 1B). A thermal plate with an open region aligning with the myobundles maintained the culture temperature at 37°C (Figure 1C). The system also included a camera-microscope unit in the CubeLab™ lid for real-time imaging of myobundle contractions and an LED array underneath the manifold for illumination. This setup enabled autonomous, in-orbit assessment of myobundle contractile function, with parallel ground controls.

**Figure 1.**
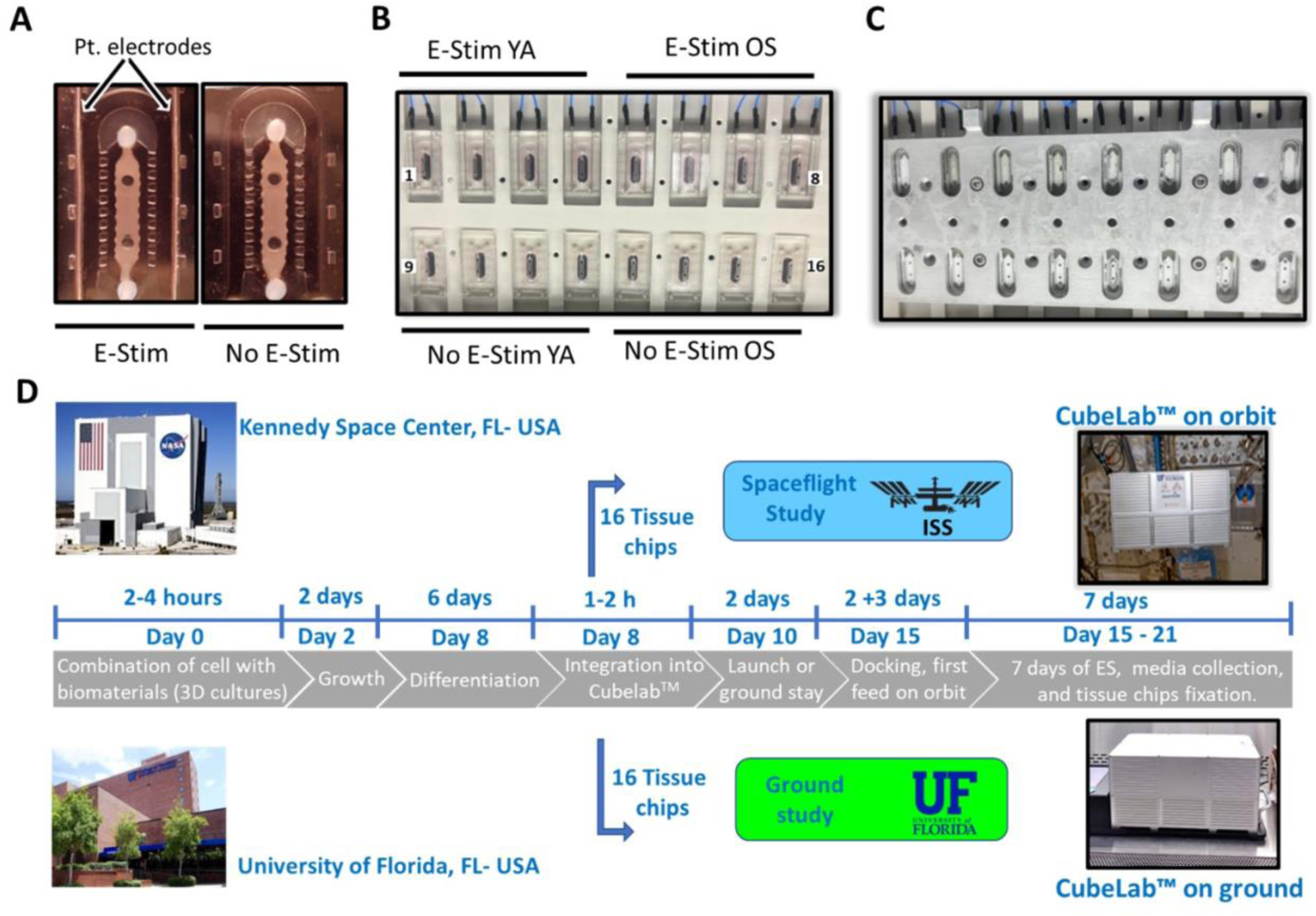
Skeletal muscle myobundle payload integration. (**A**) Skeletal muscle tissues are housed in a custom microfluidic chip with (left) or without (right) two platinum electrode leads. (**B)** Manifold layout of 16 tissue chips. The top row from left to right includes electrode stimulated young athletic chips (E-Stim YA) and old sedentary chips (E-Stim OS). The bottom row from left to right includes non-electrode stimulated young athletic chips (No E-Stim YA) and old sedentary chips (No E-Stim OS). (**C)** Frozen tissue chips at the end of the experiment are viewable through the cutout on the thermal plate. (**D)** Payload time course for the experiment performed on the ISS and asynchronously on ground. Temperature was recorded in the CubeLab^TM^ every 20 minutes except no data was collected between docking and install. Average temperature during spaceflight was 37.0 ± 0.2°C. Average temperature for the asynchronized ground study was 37.1 ± 0.1°C.

The myobundles were matured using an eight-day growth and differentiation protocol prior to launch, consistent with our previous mission (Parafati et al., 2023). Pre-flight, the tissue chips were validated for electrical stimulation responsiveness and reproducibility of peak contractile force. Myobundles in electrode-equipped chips underwent twice-daily 30-minute electrical pulses at 2 Hz for seven days, while non-electrode chips served as No E-Stim controls (Figures S1A and S1B). Contractile activity was analyzed using digital image correlation (DIC) to quantify pixel displacement within a defined region of interest (ROI), as described previously (Giza et al., 2022). YA-derived myobundles exhibited greater variability in contractile response magnitude than OS-derived myobundles, although the average displacement was not significantly different between the groups (Figure S1B).

Following protocol validation, the myobundles were prepared for spaceflight according to the experimental timeline outlined in Figure 1D. The payload launched from Kennedy Space Center and was installed on the ISS within three days. Automated protocols, including fluid exchanges every six hours for all 16 tissue chips and electrical stimulation every 12 hours for the eight electrode-equipped chips, were initiated by the Space Tango flight implementation team. After the final electrical pulse sequence, RNAlater was perfused into each chip within four hours to preserve RNA integrity, and the samples were frozen for return to Earth. A parallel ground-based study was conducted asynchronously in the same CubeLab™ hardware within a biosafety hood at the University of Florida. Experimental conditions and protocols were synchronized with the spaceflight telemetry to ensure comparability.

### Impact of donor age, E-Stim and microgravity on myofibrillogenesis

Myoblasts can be prompted to proliferate, differentiate and fuse to form mature myobundles. To compare the effects of E-Stim versus No E-Stim on myobundle formation in microgravity and on Earth, RNA was isolated post-flight from three tissue chips in each group, with RNA quantity and quality detailed in Table S1. RNA sequencing (RNA-seq) was performed on myobundles harvested on Day 21 (YA and OS, ground and flight) and compared to a reference group of myobundles formed on Day 2 (YA and OS, ground) to identify differentially expressed genes (DEGs) associated with differentiation and myogenesis under spaceflight and ground conditions (Figure 1D). The analysis revealed gene expression profiles indicative of robust muscle differentiation in both age groups (Figures S2 and S3).

Here we describe results on how e-stim, age and microgravity impact gene signatures related to differentiation, fusion and myobundle formation. In No E-Stim YA-derived myobundles, spaceflight significantly increased the number of DEGs compared to ground controls, as visualized by volcano plots (Figures S2A and S2B). Conversely, No E-Stim OS-derived myobundles exhibited similar gene expression profiles on ground and in space (Figures S2C and S2D). E-Stim YA-derived myobundles also showed a greater number of DEGs in spaceflight compared to ground (Figure S3A), while E-Stim OS-derived myobundles followed the same trend of similar response on ground and in space (Figures S3C and S3D). These findings suggest that both spaceflight and E-Stim induced a more pronounced transcriptional response in YA-derived myobundles compared to OS-derived myobundles. Gene ontology (GO) enrichment analysis across both age groups identified common pathways associated with muscle cell development and myogenesis, with shared biological processes (BP), cellular components (CC), and molecular functions (MF) across treatments (Figures S2E–H and S3E– H). GO terms such as muscle contraction, sarcomere organization, Z-disk, T-tubule, structural muscle constituents, and actin filament binding were enriched, confirming substantial transcriptional differentiation between myoblasts and myobundles regardless of E-Stim or microgravity exposure.

Canonical skeletal muscle transcripts exhibited distinct patterns across conditions. Genes promoting myofiber maturation and maintenance (e.g., *MEF2C, MYMK, MYOG*) had increased expression in Day 21 myobundles compared to Day 2 myoblasts, while early differentiation markers (e.g., *MYOD1, MYF6, MYF5*) had decreased expression (Figure 2A) (Buckingham, 1994; Francetic and Li, 2011; Klover et al., 2009; Tripathi et al., 2011; Valdez et al., 2000). Quiescence markers *PAX3* and *PAX7* showed minimal expression across conditions. Muscle-specific myosin heavy chain genes associated with type I slow-twitch fibers (e.g., *MYH3, MYH6, MYH7, MYH8*) had consistently increased expression in myobundles versus myoblasts (Figure 2B). Nebulin-related-anchoring protein (NRAP), a key component for myofibril assembly, was significantly increased in all groups (Carroll and Horowits, 2000; Lu et al., 2008), with the highest expression observed in contractile gene forming the titin complex (e.g., *TNNC3, ACTA1, TNNT3, TNNI2*) and *TCAP*, a mechanical stretch sensor (Figure 2C) (Knöll et al., 2011; Krüger and Kötter, 2016). Regulatory myosin light chains involved in Ca²⁺-dependent contractile regulation were also expressed (Figure 2D), suggesting predominant formation of type I fibers. Autocrine signaling genes (e.g., *LEP, IL6, FGF2, TGFB1*) had decreased activity across conditions (Figure 2E), whereas *IGF-2* was increased. However, *IGF-1*, promoting muscle mass, decreased, and *MSTN*, a negative regulator of muscle growth, increased in YA-derived myobundles exposed to spaceflight, suggesting an age-dependent response to microgravity (Figure 2E). Apoptosis-related genes (e.g., *BCL2, BAD, XRCC4, NOXA1*) and programmed cell death genes (e.g., *RIPK1, BAX, CASP* family) were decreased (Figure 2F). In summary, these results indicate spaceflight conditions including launch conditions support myofiber maturation and maintenance and suppressed cell death during myofibrillogenesis.

**Figure 2.**
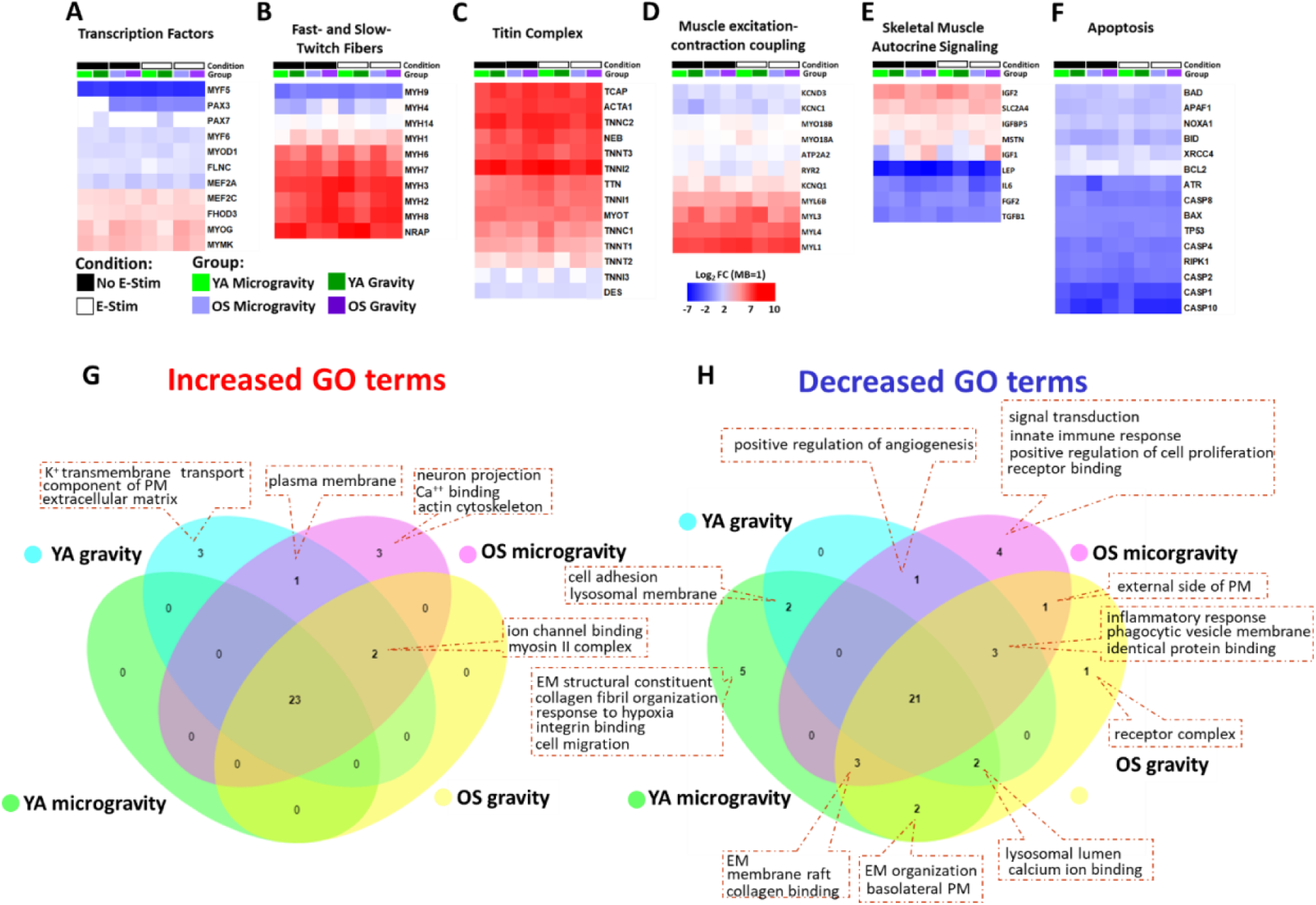
Donor-derived skeletal muscle differentiation. (**A**) Myoblast differentiation timeline for flight vs. ground control. (**B-G**) Comparisons of differential gene expression changes for No E-stim (black bars) and E-Stim (white bars) in microgravity (YA light green bars and OS light purple bars) and gravity (YA dark green bars and OS dark purple bars) as heat map log_2_ fold change (FC) mean values. Heatmaps display gene expression profiles related to categories of interest during skeletal muscle specification, based on RNA-seq data across different groups. Significance was determined according to log_2_ fold change (FC) with threshold set at ±1.5 and *p-value ≤0.05 for Day 21 myobundles vs. proliferating myoblasts at Day 2. Data are representative of three independent RNA-seq determinations. The expression variance for each gene is indicated by color key ranging from low (−7 blue) to high (+10 red). (**H-I**) Venn diagrams show common and unique GO terms based on the increased and decreased expression of DEGs (FC ≥ 1.5) identified by RNA-Seq among the comparisons of electrically stimulated YA and OS cohorts in presence or absence of gravity. GO terms identified only in the flight comparison (green and purple circles), only in the ground comparison (light blue and yellow circles), and common to both comparisons (intersections). Bonferroni corrected p-value for each GO terms was set as threshold ≤ 1e-6. Data are representative of three independent RNA-seq determinations.

GO enrichment revealed 23 increased gene activity and 21 decreased gene activity pathways shared across treatment groups (Figures 2G and 2H). Increased activity terms included muscle development and function, while decreased gene categories involved mechanosensing components and secretory pathways. Collectively, these results demonstrate that both YA- and OS-derived myobundles undergo comparable myofibrillogenesis in spaceflight and ground conditions, with an age-dependent response, notably YA-derived myobundles exhibiting a more pronounced response to microgravity, affecting muscle growth and differentiation.

### E-Stim conditioned myobundles support cellular adaptation in microgravity

Our results in Figure 2 indicate that spaceflight conditions did not significantly impact myobundle differentiation and maturation, an important finding to ensure comparability in end-point analyses. We next evaluated Day 21 myobundles cultured in microgravity versus ground conditions for both YA and OS cohorts. Global gene changes are presented as volcano plots, with pathway enrichment terms categorized into biological processes, cellular components, and molecular functions (Figures 3 and 4). In YA-derived myobundles, E-Stim induced nearly twice as many DEGs compared to No E-Stim samples predominately due to a marked increase in gene expression (1820 in E-Stim vs 646 in No E-Stim) (Figures 3A and 3B). These changes were associated with enriched pathways related to RNA binding, translation, mitochondrial function, and muscle contraction (Figures 3C and 3D). KEGG analysis identified Focal adhesion pathway (hsa04510) as the most decreased activity, while GO terms highlighted reductions in cell adhesion (GO:0007155), extracellular matrix (GO:0031012), and protein binding (GO:0005515) across both E-Stim and No E-Stim groups. These findings suggest that E-Stim partially mitigates microgravity-induced gene expression changes, although pathways related to extracellular matrix and adhesion remain susceptible to decreased expression under microgravity conditions.

**Figure 3.**
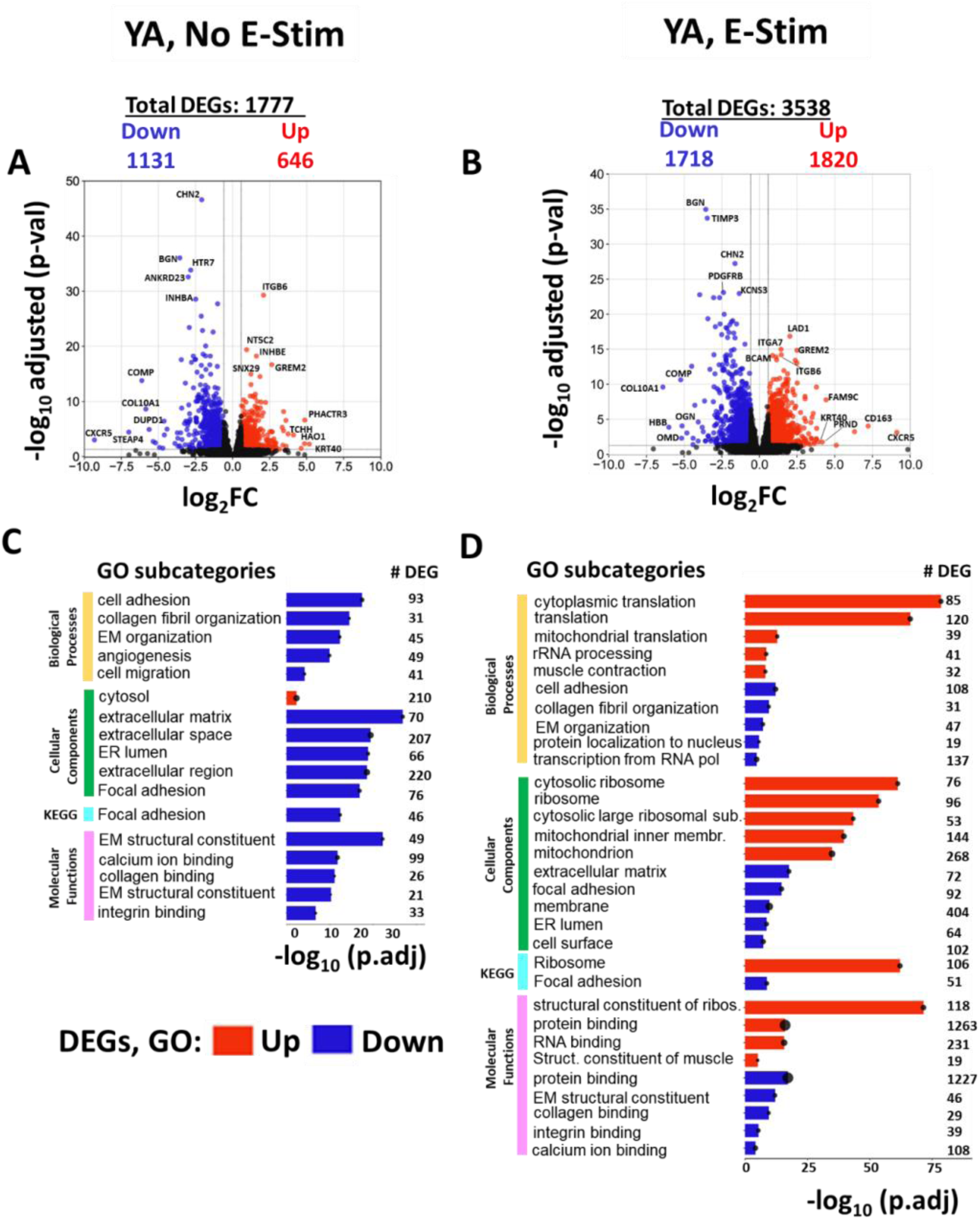
Transcriptome analysis of YA-derived myobundles exposed to microgravity vs ground. **(A, B)** Volcano plots of RNA-Seq results comparing YA-derived myobundles in microgravity vs. ground controls for (**A**) non electrically stimulated (No E-Stim) and (**B**) electrically stimulated (E-Stim) tissue chips at log_2_ fold change (log_2_FC) versus – log_10_ FDR. Red and blue circles indicated up- and down-regulation of DEGs, respectively, with at least FDR≤0.05 and log_2_FC≥±0.5 cutoff; black circles denoted non-DEGs. (**C, D**) Enriched gene ontology (GO) and KEGG analysis between microgravity and gravity control groups for (**C**) No E-Stim and (**D**) E-Stim tissue chips. Pathway enrichment analysis by biological processes (yellow), cellular component (green) and molecular function (pink), and Kyoto Encyclopedia of Genes and Genomes (KEGG, light blue) was performed using the Database for Annotation, Visualization, and Integrated Discovery (DAVID) gene functional classification tool on the increased and decreased expression of DEGs (FDR≤0.05 and log_2_FC≥± 0.5) between the indicated groups based on a two-tailed Student’s t test. Bar dots were ordered according to the significance and are ranked by log_10_ FDR adjusted p-values. The horizontal axis represents the log_10_ adjusted p-values of enriched DEGs to background genes, while the vertical axis represents the enriched subcategories name. Numbers in parenthesis indicate the number of enriched DEGs reported in each sub-ontology category. Data are representative of three independent RNA-seq determinations.

**Figure 4.**
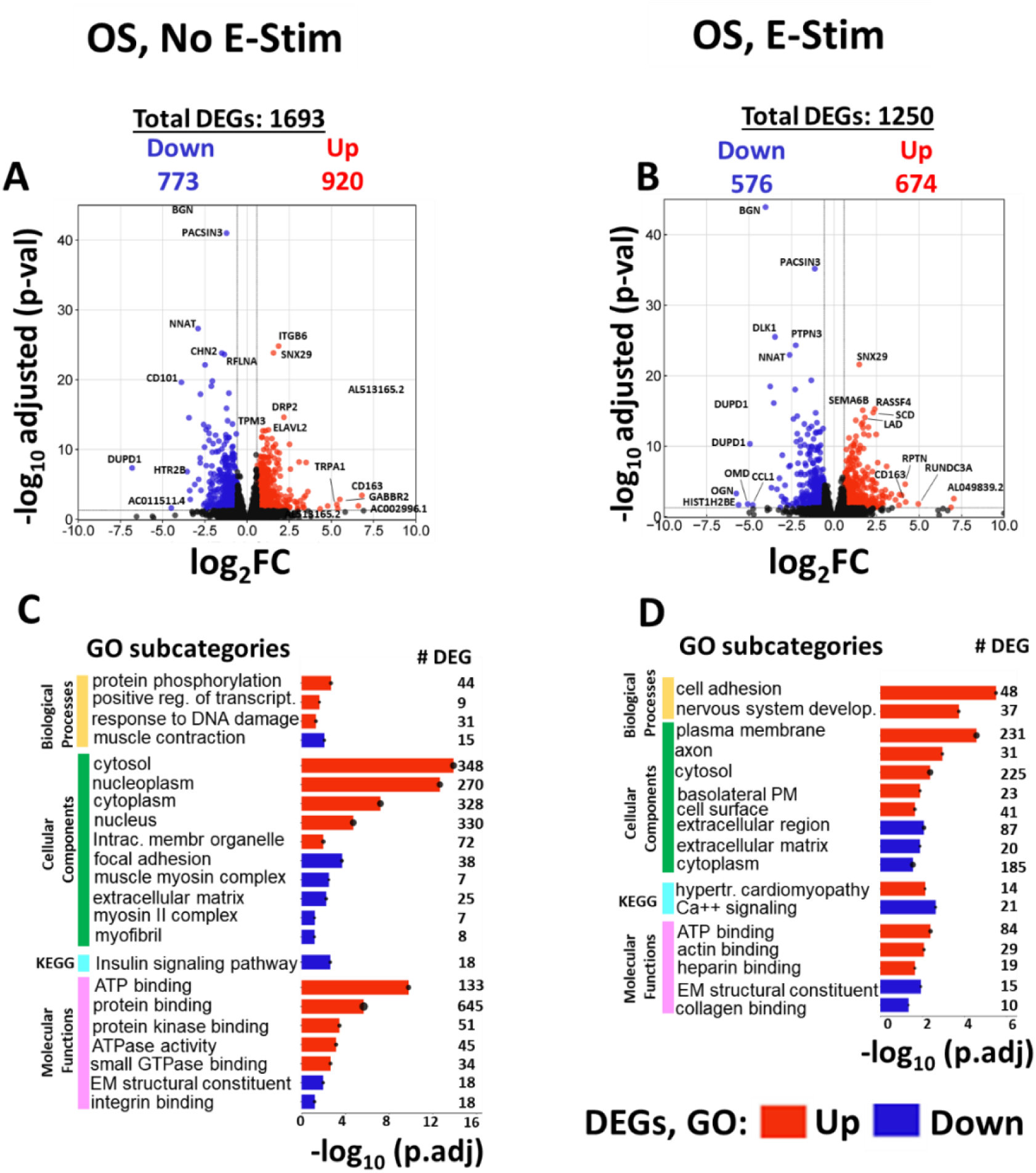
Transcriptome analysis of OS-derived myobundles exposed to microgravity vs ground. **(A, B)** Volcano plots of RNA-Seq results comparing OS-derived myobundles in microgravity vs. ground controls for (**A**) non electrically stimulated (No E-Stim) and (**B**) electrically stimulated (E-Stim) tissue chips at log_2_ fold change (log_2_FC) versus – log_10_ FDR. Red and blue circles indicated up- and down-regulation of DEGs, respectively, with at least FDR ≤ 0.05 and log_2_FC ≥ 0.5 cutoff; black circles denoted non-DEGs. (**C, D**) Enriched gene ontology (GO) and KEGG analysis between microgravity and gravity control groups for (**C**) No E-Stim and (**D**) E-Stim tissue chips. Pathway enrichment analysis by biological processes (yellow), cellular component (green) and molecular function (pink), and Kyoto Encyclopedia of Genes and Genomes (KEGG, light blue) was performed using the Database for Annotation, Visualization, and Integrated Discovery (DAVID) gene functional classification tool on the up- and down-regulated DEGs (FDR ≤ 0.05 and log_2_FC ≥ 0.5) between the indicated groups based on a two-tailed Student’s t test. Bar dots were ordered according to the significance and are ranked by log_10_ FDR adjusted p-values. The horizontal axis represents the log_10_ adjusted p-values of enriched DEGs to background genes, while the vertical axis represents the enriched subcategories name. Numbers in parenthesis indicate the number of enriched DEGs reported in each sub-ontology category. Data are representative of three independent RNA-seq determinations.

OS-derived myobundles displayed a contrasting pattern: E-Stim under microgravity reduced the total number of DEGs compared to No E-Stim in contrast to the increased DEGs observed in the YA cohort (Figures 4A and 4B). Increased DEGs in both E-Stim and No E-Stim OS groups were associated with cytosol (GO:0005829) and ATP binding (GO:0005524). However, cell adhesion (GO:0007155) was exclusively enriched in E-Stim group only (Figures 4C and 4D). KEGG analysis identified the hypertrophic cardiomyopathy pathway (hsa05410) as enriched in the E-Stim group (Figure 4D). Decreased DEGs in OS-derived myobundles were linked to extracellular region (GO:0005576) and extracellular matrix structural constituents (GO:0005201). Muscle contraction (GO:0006936) and muscle myosin complex pathways (GO:0005859, GO:0016460) were uniquely decreased in No E-Stim samples but remained unaffected in the E-Stim group. KEGG pathway analysis also revealed downregulation of the insulin signaling (hsa04910) in No E-Stim samples and downregulation of the calcium signaling pathway (hsa04020) in E-Stim samples (Figures 4C and 4D).

These results suggest that E-Stim supports adaptation to microgravity in OS-derived myobundles by mitigating transcriptional response associated with DNA damage response and protein phosphorylation compared to No E-Stim samples. Furthermore, E-Stim appears to promote transcriptional programs linked to cell adhesion and plasma membrane dynamics that support muscle function, particularly in the OS cohort. These findings underscore the differential impact of E-Stim on gene expression in YA-versus OS-derived myobundles under microgravity, with E-Stim showing a potential protective effect against microgravity-induced cellular dysfunction.

### Mitochondrial pathways are altered in E-Stim YA and OS myobundles in microgravity

The impacts of microgravity on mitochondrial function have been well-documented in prior studies (Bonanni et al., 2023; Corydon et al., 2023; Da Silveira et al., 2020; Feger et al., 2016; Nguyen et al., 2021). Our RNA sequencing analysis revealed significant changes in gene expression related to mitochondrial energetics under microgravity, with distinct responses observed between YA- and OS-derived myobundles. In the YA cohort subjected to E-Stim, 66 DEGs related to mitochondrial function were increased. These included genes involved in proton-transporting ATP synthase (e.g., *ATP5F1D, ATP5PD, ATP5PB, ATP5F1C, ATP5MF, ATP5MG*), mitochondrial electron transport (e.g., *UQCRH, UQCR11, UQCRQ, UQCRB*), cytochrome c oxidase (e.g., *COX7B, COX5B, COX6B1, COX7A2L, COX5A, COX7A1*), NADH dehydrogenase/ubiquinone (e.g., *NDUFA10, NDUFB9, NDUFA4, NDUFS5, NDUFV1, NDUFA9, NDUFB1, NDUFA2, NDUFS7, NDUFB7, NDUFS8*), and outer mitochondrial membrane organization (e.g., *TOMM7, TOMM20, TOMM22*). Conversely, decreased DEGs were primarily linked to the PI3K signaling pathway, which stimulates cytoskeletal arrangement and cell survival (e.g., *PIK3CA, PIK3CB, PIK3C2A, PIK3CD, PIK3R2, PIK3R1*) (Figure 5A).

**Figure 5.**
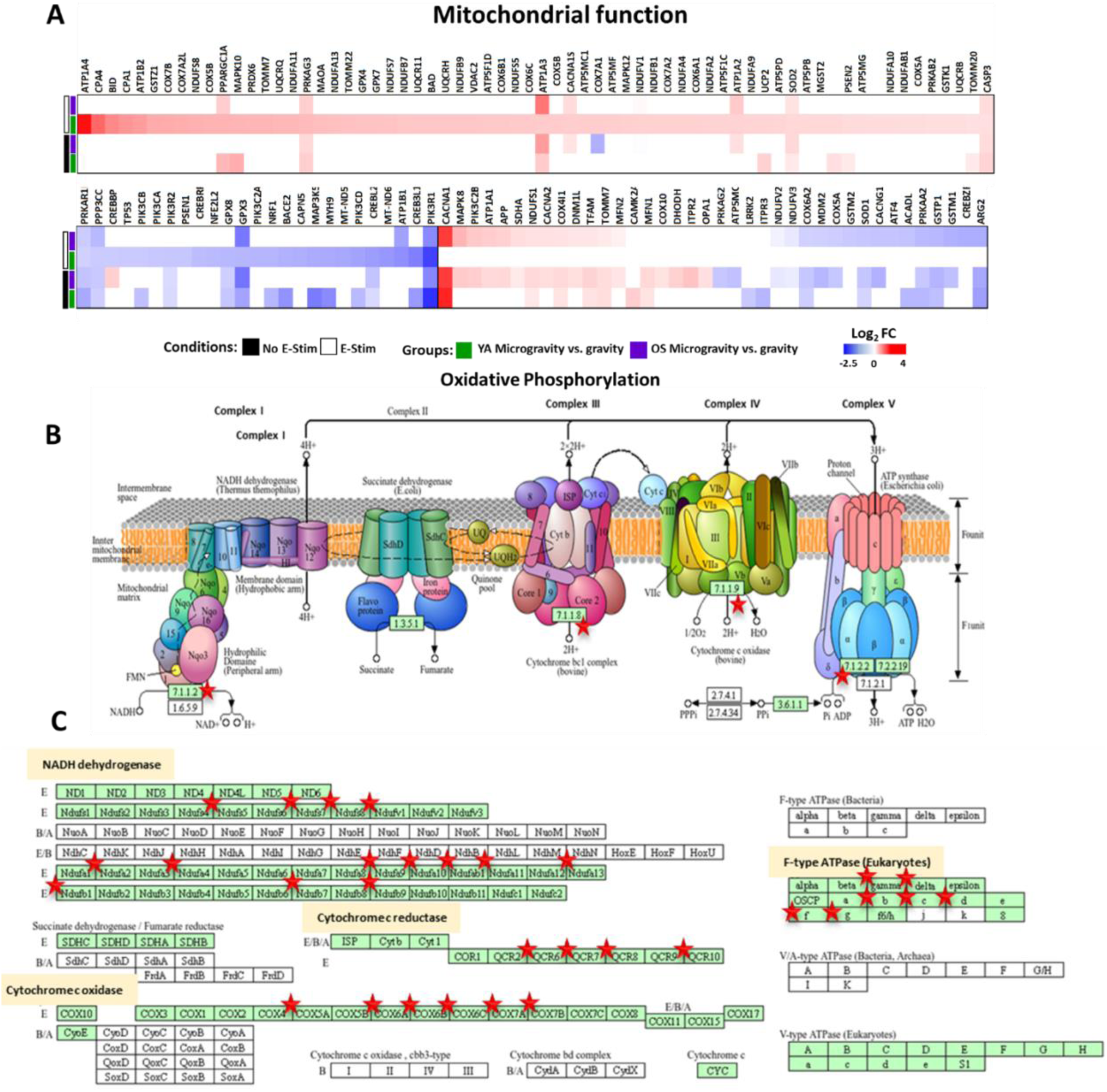
YA E-Stim group shows genes with increased activity involved in oxidative phosphorylation under microgravity vs gravity. **(A)** Heatmap of DEGs related to mitochondrial function and their expression intensity across the treatment groups under microgravity vs gravity. (**B**) Respiratory chain complexes involved in oxidative phosphorylation. This pathway was identified as the most significantly enriched pathway in a KEGG pathway analysis, highlighting transcripts uniquely expressed in the E-Stim YA microgravity group compared to gravity. (**C**) Subunits of each respiratory chain complex displayed as boxes. The DEGs highlighted by a red star indicate those genes with increased activity in response to microgravity. Significance was determined according to log_2_FC with threshold set at ± 0.5 and FDR ≤ 0.05 vs. gravity. Data are representative of three independent RNA-Seq determinations.

In contrast, the OS cohort exhibited upregulation of 20 DEGs regardless of E-Stim treatment. These genes encode for proteins involved in mitochondrial fusion and important for mitochondrial dynamics and quality control (e.g., *OPA1, NDUFS1, DHODH, COX10, APP, SDHA, COX4I1, MFN1, MFN2*), voltage-gated calcium channels (e.g., *CACNA2D1, CACNA1E, ITPR2*), nucleotide binding and ion transport (e.g., *ATP1A1, ATP1A2, ATP1A3, PIK3C2B, MAPK8, DNM1L*), and reactive oxygen species (ROS) regulation (*SOD2*) (Figure 5A). However, decreased DEGs were associated with antioxidant and mitochondrial enzyme activities including glutathione transfer (e.g., *GSTP1, GSTM2, GSTM1*), superoxide dismutase (SOD1), NADH dehydrogenase (e.g., *NDUFV2, NDUFV3*), and cytochrome oxidase (e.g., *COX5A, COX6A2*). Notably, activating transcription factor 4 (*ATF4*), a gene associated with muscle atrophy(Adams et al., 2017), was uniquely decreased in E-Stim OS-derived myobundles, suggesting a protective effect against muscle degradation in microgravity.

Our analysis highlighted a strong signature of mitochondrial energetics in the E-Stim YA-derived myobundles under microgravity compared to ground controls. As depicted in Figure 5B, the expression of genes encoding components of the respiratory chain complexes involved in oxidative phosphorylation was increased. These DEGs included subunits of Complex I (mitochondrial encoded NADH dehydrogenase), Complex III, Complex IV (cytochrome c oxidase subunits) and Complex V (ATP synthase). These results underscore the critical role of mitochondrial respiratory chain complexes in adapting to microgravity conditions in E-Stim treated YA-derived myobundles (Figure 5C).

### E-Stim enhances fiber type adaptations and mitochondrial biogenesis in microgravity

We evaluated the contractile response of muscle tissue chips to 30 min of E-Stim applied every 12 hr for seven days by recording videos before and during delivery of low frequency electrical pulses through the platinum electrodes. Representative images of YA- and OS-derived myobundles in microgravity and on Earth are shown in Figure 6A, captured inside the CubeLab^TM^. Contractile magnitude was determined by analyzing 40-second video sequences captured at 40 frames per second during E-Stim relative to a reference image taken prior to stimulation. Functional assessments were based on the average contractile response of multiple E-Stim events per tissue chip over seven days. Displacement measurements plotted over time displayed two peaks per second for electrically synchronized muscle contractions stimulated at 2 Hz frequency (Figure 6B). On Earth, YA-derived myobundles exhibited higher contractile magnitude (10.4 ± 3.2 μm) compared to OS-derived myobundles (5.6 ± 1.9 μm) (Figure 6C). CubeLab^TM^ data aligned closely with results from ground-based incubator experiments (Figure S1), with YA-derived myobundles demonstrating greater variability in contractile magnitude compared to the older cohort. In microgravity, contractile function declined, with average contractile magnitudes of 7.7 ± 4.6 μm for YA-derived myobundles and 4.7 ± 1.0 μm for OS-derived myobundles (Figure 6C). The difference in contractile magnitude values between YA and OS on ground was statistically different, however, the values were not significantly different in microgravity indicating a decline in function in microgravity.

**Figure 6.**
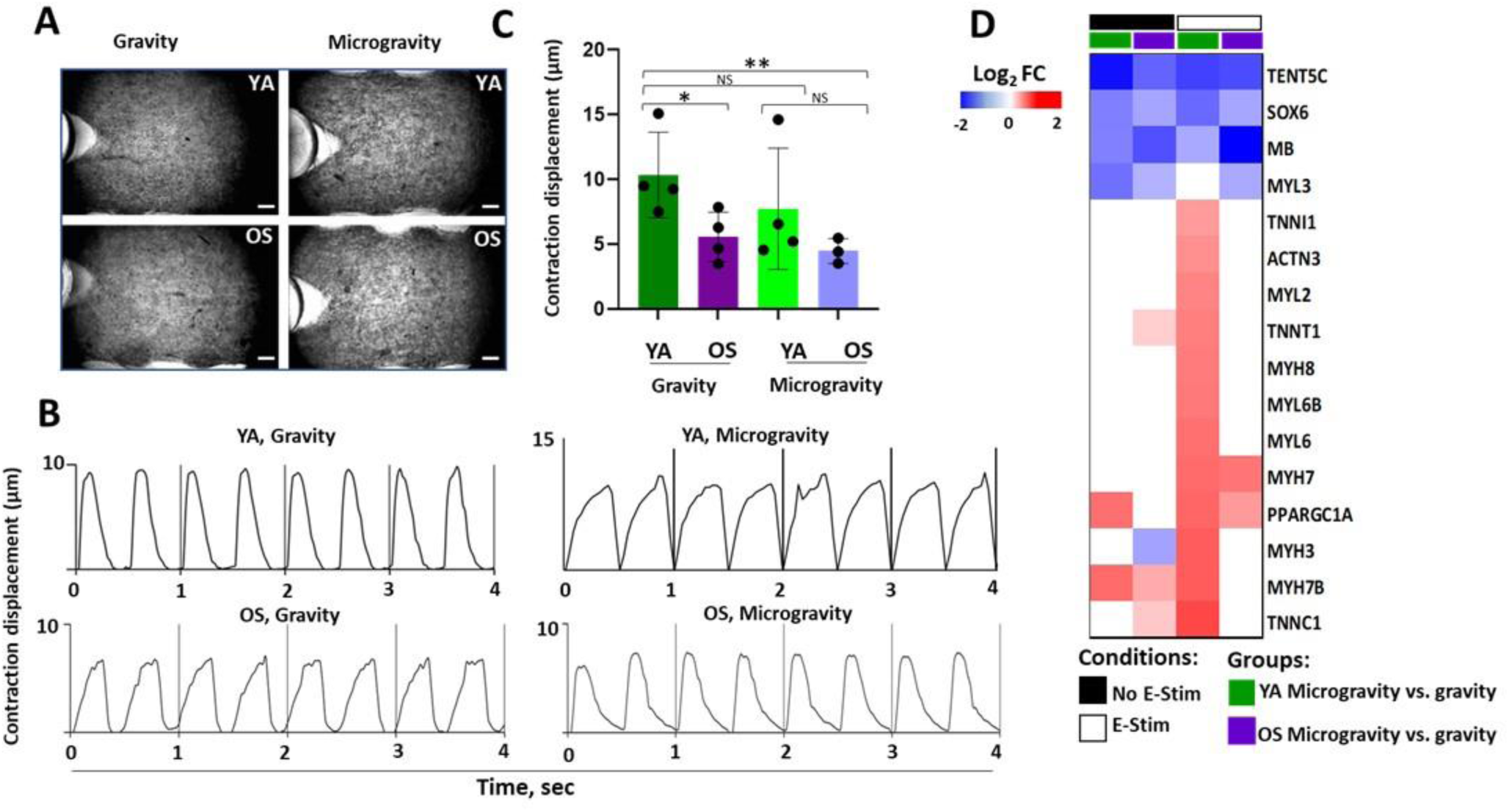
The effect of microgravity on contraction magnitude and fiber type in donor-derived myobundles. **(A)** 10 x representative photographs (size: 3088 × 2076 pixel; 2.6 x 1.7 mm, scale bar 200um) of donor-derived myobundles in gravity and microgravity conditions (**B**) Contraction displacement values (µm) between young and old myobundles cultured for 12 days. E-Stim regime (voltage: 3 V, frequency: 2 Hz, duration: 2 ms) was applied for 30 min with 12 h rest for 7 days. Video was recorded for 40 s at 40 fps. As the control, donor-derived myobundles were cultured without E-Stim with contraction displacement values close to zero. Each data point is average of two recordings per tissue chip. Statistical significance was calculated using the two-tailed, unpaired Student’s t-test: YA gravity vs. OS gravity, *p≤0.046; YA gravity vs. OS microgravity, **p≤0.032; YA gravity vs. YA microgravity and YA microgravity vs. OS microgravity, NS (not significant), p>0.05. (**C**) Heatmap of expression levels for muscle fiber type I genes between microgravity vs. gravity in absence or presence of E-Stim. Significance was determined according to log2FC mean values with threshold set at ±0.5 and FDR≤0.05 vs. gravity. The expression variance for each gene is indicated by color key ranging from low (blue) to high (red). White denotes no changes in gene expression. Data are representative of three independent RNA-seq. determinations.

A heatmap analysis of muscle fiber type-associated gene expression revealed increased expression of genes encoding myosin light and heavy chains, as well as troponin isoforms (e.g., *MYL2, MYH6, MYH7, MYH3, MYH8, TNNI1, TNNT1, TNNC1, MYH7B*) in E-Stim YA-derived myobundles under microgravity compared to ground controls (Figure 6D). This expression pattern is consistent with type 1, slow-twitch muscle phenotype. Conversely, *ACTN3*, a marker of fast-twitch muscle fibers, was also increased, suggesting a mixed fiber type composition. This mixed fiber profile was not observed in the No E-Stim group. For OS-derived myobundles, E-Stim promoted the upregulation of *MYH7*, indicative of slow-twitch muscle adaptations in microgravity. Furthermore, sarcomeric genes such as *MYH7B, TNNT1*, *TNNC1* had increased expression in the No E-Stim OS-derived group, indicating slow-twitch muscle adaptations even in the absence of E-Stim. We also observed significant increased expression of *PGC-1α*, a transcriptional co-activator that promotes mitochondrial biogenesis and oxidative metabolism, in YA-derived myobundles, regardless of E-Stim treatment or No E-Stim under microgravity conditions versus ground. *PGC-1α* plays a central role in promoting type I muscle fiber phenotype (Lin et al., 2002). Conversely, *SOX6*, a transcriptional suppressor of slow-twitch muscle fiber genes, had consistently decreased activity across treatment groups. This is consistent with previous findings showing that mice lacking *SOX6* display an increased proportion of slow-twitch muscle fibers (An et al., 2011; Song et al., 2022). These results suggest that E-Stim enhances slow-twitch muscle adaptations and mitochondrial biogenesis in both YA- and OS-derived myobundles, particularly under microgravity conditions.

### Microgravity reveals molecular pathways affected in young and aged muscle phenotypes

We compared gene expression profiles across the four treatment groups cultured in the same CubeLab^TM^ environment under microgravity and ground conditions to identify unique genes influenced by microgravity exposure (Zhao et al., 2019). The dataset is visualized as a circos plot, illustrating the overlap of shared genes (purple lines) and functional terms (light blue lines) between microgravity and ground conditions (Figure 7A). Among the cohorts, the E-Stim YA group displayed the highest number of DEGs, with 26% of functional terms overlapping across the dataset (471 of 1820; Figure 7A). In contrast, other cohorts exhibited fewer DEGs but a higher degree of overlap, exceeding 50%. This suggests that the E-Stim YA group undergoes the most extensive transcriptional changes.

**Figure 7.**
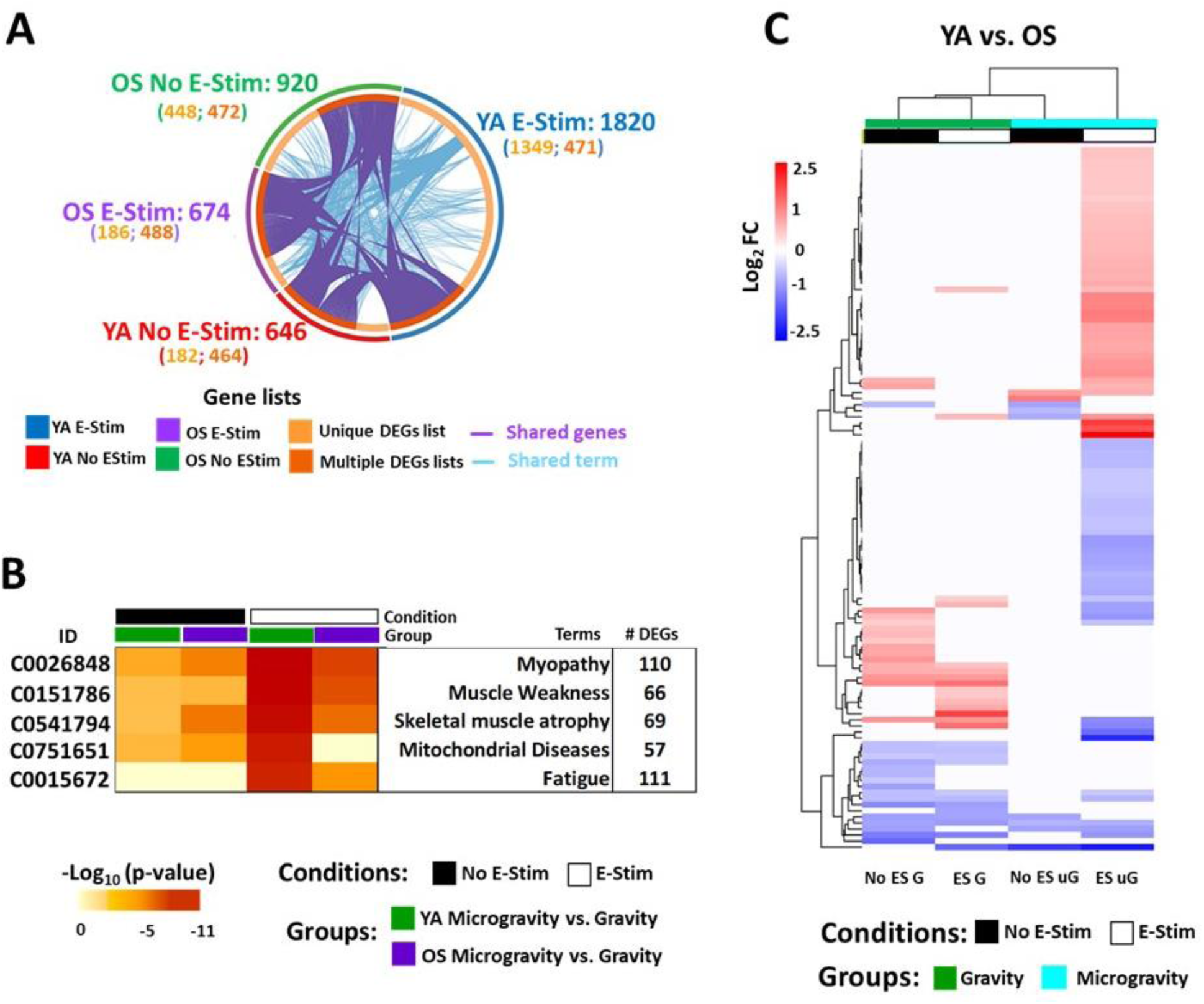
Functional enrichment meta-analysis highlights muscle atrophy pathways and modulation of muscle aging genes in microgravity. (**A**) Circular plot depicts how genes (purple lines) and functional terms (light blue lines) from the different input gene lists of genes with increased activity overlapping under microgravity vs gravity. The outer circle represents the identity of each gene list (YA E-Stim, blue; YA No E-Stim, red; OS E-Stim, purple; OS No E-Stim, green). The inner circle represents shared genes (dark orange) and unique genes (light orange). The number of genes in these two categories is indicated in the parenthesis. (**B**) Heatmap of the top five terms driving the clustered dataset. Pathways are sorted by -Log10 p-values. Light orange to dark orange indicates low to high value, respectively. (C) Hierarchical clustering analysis of muscle aging genes differentially expressed as log_2_FC mean values between YA vs OS myobundles in gravity and microgravity when compared with the muscle aging reference list (Su et al., 2015). Each column contains expression values for YA vs. OS in absence and presence of E-Stim. Significance was determined according to log_2_FC with threshold set at ± 0.5 and FDR≤ 0.05. The legend indicates the relation between log_2_FC-scaled expression values and colors, and the colors were balanced. White represents no significative changes in gene expression. Data are representative of three independent RNA-seq. determinations.

To investigate gene-disease associations, a meta-analysis of genes with increased activity was performed using the DisGeNET database (Figure 7B) (Piñero et al., 2020). The analysis identified multiple dysregulated disease ontology terms related to muscle dysfunction in the tissue chip exposed to microgravity compared to ground controls. A heatmap of DisGeNET muscle disease-related pathways highlights the most enriched pathways, ranked by log10 p-values, with a DEG threshold >55 (Figure 7B). The most significant disease terms associated with the combined dataset, driven by the YA E-Stim cohort, included myopathy (C0026848, 110 DEGs), muscle weakness (C0151786, 76 DEGs), skeletal muscle atrophy (C0541794, 69 DEGs), mitochondrial diseases (C0751651, 57 DEGs) and fatigue (C0015672, 111 DEGs). By mining DEGs from DisGeNET muscle disease-related pathways with the Rummagene tool, 61 genes overlapping with most significant myopathy term (C0026848) were identified (Figure 7B) (Clarke et al., 2024). These findings link microgravity exposure to muscle atrophy in human tissue chips, corroborating previous studies (Juhl et al., 2021; Kim et al., 2024; Oommen et al., 2024).

Finally, we directly compared the YA versus the OS groups under microgravity and ground conditions to assess age-related transcriptional differences. Using a database of muscle-aging genes derived from a meta-analysis of 2852 public gene expression arrays in skeletal muscle(Su et al., 2015), we identified genes affected by microgravity by cross-referencing 484 genes increased in human aged skeletal muscle with our dataset. Differential expression analysis between the YA vs OS groups is presented as a hierarchical clustering map (Figure 7C). The E-Stim YA vs E-Stim OS comparison in microgravity exhibited the greatest change in expression. 45 aging genes were significantly increased while a further 41 aging genes were significantly decreased. These genes were not changed when comparing No E-Stim YA to No E-Stim OS under microgravity nor was genes expression altered when comparing the cohorts on ground.

In electrically stimulated YA vs OS cohorts under microgravity, we observed an increase in aging genes related to inflammation and immune response (e.g., *ALOX5AP, CCR10, CFD*), cell signaling and growth (e.g. *WISP2, IRS2, SIRT1*), RNA processing (e.g. *CLK4, IFRD1, DUSP1*), cellular structure and cytoskeleton regulation (e.g. ABRA, AHNAK, MYL6B), and metabolism and stress response (e.g., ASS1, SAT1, AKR1C3, TRMT112) (Table S2). *ALOX5AP* expression, encodes for 5-lipoxygenase-activating protein crucial for activating Alox5, increased by 2.5-fold in E-Stim YA vs E-Stim OS under microgravity. This protein has been shown to be increased in aged human skeletal muscle and following glucocorticoid treatment(Kim et al., 2022). However, increased expression of *WISP2, IRS2, SIRT1* suggests enhanced muscle regeneration, improved insulin sensitivity and resistance to oxidative stress and adaptive response in the electrically stimulated YA cohort under microgravity (Kang et al., 2023; Powers and Schrager, 2022; Yuan et al., 2016).

Our data revealed a decrease in aging gene expression related to inflammation and immune response (e.g, *TLR4, SERPING1, LAMP1*), autophagy (e.g., *SIDT2, TECPR2*), cellular stress and senescence (e.g. *TRIM8, WARS, AKAP12*) and extracellular matrix remodeling (e.g., *ADAMTS5, SPOCK1*). We observed a decrease in FOXO1, A key transcription factor in the insulin/IGF-1 signaling pathway, may indicate suppression of muscle protein breakdown during E-Stim. Furthermore, we observed a 2-fold decrease in toll-like receptor 4 (*TLR4*) expression. Previous studies have shown that stimulated myotube contractions reduce membrane-bound TLR4 while increasing soluble TLR4, thereby inhibiting TLR4-mediated signaling and preventing myotube atrophy (Ducharme et al., 2023).

These findings highlight significant transcriptional differences between young and old muscle phenotypes under microgravity, providing insights into how microgravity influences or accelerates age-related changes in skeletal muscle and highlight the potential role of electrical stimulation in partially mitigating tissue degradation.

## DISCUSSION

Understanding the impact of microgravity on muscle cell biology is critical to developing therapeutic countermeasures to prevent deconditioning and accelerated aging of skeletal muscle during spaceflight. Our study employed morphological and transcriptomic approaches to investigate adaptation of donor-derived skeletal muscle myobundles to microgravity. We utilized Space Tango’s CubeLab™ for controlled and autonomous in vitro experiments and evaluated the effects of microgravity on myobundle contractile function and cellular mechanisms through global transcriptome profiling. Our muscle lab-on-chip approach enabled direct comparison between eight young- and eight older-derived myobundles in microgravity compared to gravity, highlighting decline in contractile displacement magnitude, enrichment of gene pathways associated with muscle weakness, and alteration of age-associated gene expression in microgravity. As we hypothesized, the younger donor-derived myobundles exhibited more pronounced transcriptional adaptations to microgravity particularly associated with mitochondrial oxidative phosphorylation compared to myobundles derived from older donors. Our findings underscore the utility of 3D muscle tissue chips for studying muscle biology adaptations in space. As most studies investigating the impact of microgravity on muscle atrophy have been conducted in tissues isolated from humans and rodents in space, our system offers the unique opportunity to interrogate the muscle cell’s autonomous response to microgravity without the influence of tissue innervation, circulation, and immune cell interactions.

Gene Ontology analysis revealed a 55% similarity between young and old donor-derived myobundles, reflecting a coordinated transition from proliferation to skeletal muscle-specific functions, including intracellular trafficking and extracellular matrix remodeling. On the other hand, microgravity uniquely influenced specific pathways during myogenesis. In young, space-flown myobundles, pathways related to extracellular matrix organization, hypoxia response, and integrin binding had decreased activity, indicating diminished intracellular signaling and muscle maintenance. Conversely, old space-flown myobundles exhibited an increased level of activity in pathways associated with calcium binding, neuron projection, and cytoskeletal function, suggesting compensatory mechanisms to mitigate microgravity-induced atrophy by enhancing neurotransmitter sensitivity. KEGG pathway analysis revealed disruptions in striated muscle pathways, such as cardiac contraction, and alterations in phagosome-lysosome dynamics, which as critical for cellular homeostasis. These results suggest that microgravity disturbs the balance between intracellular homeostasis and external forces, resulting in changes to the cytoskeletal, signaling, and membrane permeability changes during myogenesis.

Comparative analysis between microgravity and ground control conditions on Day 21 revealed DEGs in young and old myobundles, especially under electrical stimulation. Young myobundles without stimulation exhibited significant decreased activity of genes involved in cell adhesion, extracellular matrix organization, and calcium ion binding, processes essential for muscle function and repair (Mahmassani et al., 2017). Genes with decreased activity included cadherin 11, glycoprotein thrombospondin 1, and collagen 10α1, emphasizing microgravity’s adverse effects on cellular matrix integrity and muscle health (Ricci et al., 2021; Shinde et al., 2016). Similar reductions in focal adhesion formation have been observed in simulated microgravity studies (Guignandon et al., 2003; Tan et al., 2018). These findings are important as the extracellular matrix plays a critical role in muscle fiber force transmission, maintenance, and repair.

Young myobundles showed adaptive responses, such as increased expression of mitochondrial and translational genes (e.g., cytochrome C oxidase subunit 4I1) to support ATP production, despite potential oxidative stress and reactive oxygen species (ROS)-induced muscle atrophy (Lee et al., 2022; Zhao et al., 2019). Microgravity also triggered stress-related pathways, including upregulation of eukaryotic translation elongation factor 1 alpha and GADD45G, indicating cellular recovery mechanisms, although GADD45G may also contribute to atrophy (Yoshida and Delafontaine, 2020). In older myobundles, insulin-like growth factor-1, critical for muscle maintenance, had decreased expression, potentially reflecting impaired adaptive capacity (Ascenzi et al., 2019; Yoshida and Delafontaine, 2020), while upregulation of tribbles pseudokinase 3 and myostatin (MSTN) reflected endoplasmic reticulum stress and muscle atrophy, consistent with prior findings linking MSTN inhibition to prevention of microgravity-induced muscle loss (Gao et al., 2018; Kato and Du, 2007; Lee et al., 2022; Ren et al., 2024).

Overall, young myobundles exhibited coordinated stress responses and adaptations to microgravity, while older myobundles showed reduced responsiveness. These results highlight the complex interplay between microgravity and skeletal muscle biology, emphasizing the need for further studies to validate these mechanisms and their potential implications for muscle health in space. Previous reports have documented reduced skeletal muscle mass and contractile force in microgravity in primate and rodent models (Chopard et al., 2000; Ren et al., 2024; Stein, 2013), with type II fiber atrophy resembling sarcopenia (Gao et al., 2018; Nilwik et al., 2013). Our study extends these findings by showing muscle cell autonomous structural and functional deficits in old-donor myoblasts and reduced twitch magnitude under simulated microgravity (Giza et al., 2022; Ren et al., 2024). Transcriptomic analysis revealed elevated expression of slow-twitch (type I) fiber-related genes (e.g. *MYH3*, *MYH7*) and sarcomeric components (e.g. *MYL6*, *TNNI1*) as compensatory adaptations to counteract muscle loss.

A meta-analysis of four independent RNA-Seq datasets identified significant gene signatures linked to muscle disease pathways, reflecting physiological responses to reduced muscle load imposed by microgravity. DEGs associated with myopathy encoded proteins essential for maintaining skeletal muscle mass and function, establishing a link between microgravity, altered gene expression, and muscle atrophy (Juhl et al., 2021; Kim et al., 2024; Oommen et al., 2024). Furthermore, we observed gene expression and functional changes induced by microgravity resembling those seen in muscle aging, such as reduced contractility and downregulation of cell adhesion and ECM binding genes (Schüler et al., 2021; Wood et al., 2014). Enhanced expression of mitochondrial genes necessary for ATP production, coupled with increased ROS, suggests a dynamic response that may drive oxidative stress and activate downstream cellular processes (Espinosa-Jeffrey et al., 2016; Forghani et al., 2024).

Performing experiments in an autonomous CubeLab^TM^ has the advantage of minimizing crew time and advancing miniaturization and turnkey technologies for evaluating cell responses to different stimuli in normal and extreme environments and over long durations. This approach also has limitations. Sample size is limited by the volume of media that can be stored in the CubeLab^TM^. In our system, 16 tissue chips underwent perfusion, imaging, electrical stimulation and video capture. Due to programing and data storage capacity, only one of two electrical stimulation events were recorded per chip per 24 hr and downlinked. Maintaining an ideal cell incubator environment with high humidity levels in a closed electronic system also poses some challenges. Bubble traps were installed to successfully eliminate air in the tissue chips. Overall, iterative access to the ISS enabled our team to implement important advancements in tissue chip technology from our previous mission (Parafati et al., 2023).

Our study provides an integrated gene- and network-level meta-analysis, identifying key molecular signatures and candidate driver genes involved in muscle cell adaptation to microgravity, with a focus on age and electrical stimulation. We highlighted mediators such as extracellular matrix remodeling factors, mitochondrial markers, and senescence indicators, providing new insights into mechanisms underlying microgravity-induced muscle alterations and disease progression. Notably, inefficient muscle adaptation to microgravity may exacerbate inflammation and potentially accelerate the onset of muscle wasting (Kunz and Lanza, 2023; Parafati et al., 2023). Upregulation of chemokine signaling-related genes in microgravity suggests tissue inflammation, with chemokines known to be induced in dystrophic muscle (De Paepe and De Bleecker, 2013; De Paepe et al., 2012; Demoule et al., 2005). A decrease in genes associated with inflammation, autophagy, and cellular senescence and in particular FOXO1 and TLR4 in electrically stimulated young cohort compared to the older electrically stimulated cohort suggests a role for electrical stimulation in reducing tissue degradation. Circulating biomarkers, such as extracellular vesicles, identified as potential indicators of inflammation-related muscle changes (Lombardo et al., 2024), have potential to serve as non-invasive diagnostic tools to monitor muscle-specific changes in microgravity. These findings could facilitate the development of targeted interventions to safeguard astronaut health and provide biomarkers for disease progression to benefit health on Earth.

## METHODS

### Patients and Muscle Biopsy Data Analysis

For this study, we used frozen muscles biopsies obtained from the middle region of the vastus lateralis of five young, active white male patients (mean 35 ± 4) and five old, sedentary white male patients (mean 70 ± 4) with no known mutations (AdventHealth ID) as described (Sparks et al., 2011). Participants were considered active if they engaged in endurance exercise at least 3 days a week without extensive lay off over the previous 6 months. Participants were considered sedentary if they completed one or fewer structured exercise sessions a week. All patients have given informed written consent. and the study protocol was reviewed and approved by the institutional review board at AdventHealth, Orlando (IRBNet #554559). Thus, the study was performed in accordance with the ethical standards laid down in the1964 Declaration of Helsinki. Satellite cells were isolated from the biopsies (quiescent mononuclear muscle cells) by trypsin digestion and pre-plated as described (Sparks et al., 2011). Samples were obtained sterile and free of myoplasma at passage p2. Muscle biopsy data for donor-derived primary skeletal muscle were previously published (Giza et al., 2022).

### Myoblast purification

To avoid contamination by non-muscle cells, mainly fibroblasts, harvested cells were purified with an immuno-magnetic sorting system (Miltenyi Biotec, Paris, France) using an anti-CD56/NCAM antibody (Giza et al 2022). Purified myoblasts were plated in collagen-coated Petri dishes and cultured in skeletal muscle cell growth medium (Promocell Cat# C-23060) supplemented with 0.1 mg/ml primosin to protect cultures from microbial contamination, 5% FBS and gentamicin at 37°C in humidified atmosphere with 5% CO_2_. At cell isolation, all human myoblasts were at passage 4 (p4) and expanded no more than twice. Passage numbers were matched for young- and old-derived patients’ cells. All experiments were performed at p6, to avoid premature replicative senescence of myoblasts CD56+, which is known to interfere with proliferation and differentiation processes. Myoblasts were kept from reaching confluency to avoid differentiation prior to pooling cells in 1:1 ratio and cryopreserving aliquots. Enrichment was confirmed by comparing CD56^−^ and CD56^+^ cell populations by FACS staining as described (Giza et al., 2022).

### Myobundle Culture and Maintenance in CubeLab^TM^

Donor-derived 3D myobundles were generated by seeding myoblasts-derived from a pool of two cohorts (young and old) in tissue-engineered constructs and launched to the ISS aboard during the SpaceX CRS-25 commercial resupply service mission for two weeks (**Figure 1**). We developed young, active (YA) and old, sedentary (OS) miniaturized fused 3D myotube culture chip made in polydimethylsiloxane (PDMS) with contraction monitoring capacity by culturing the myoblasts in differentiation and maintenance media for a total of 21 days and electrically stimulated the last seven days. Cell seeding was performed at NASA’s Kennedy Space Station Processing Facility (SSPF) for flight and at the University of Florida laboratory for the ground test. Injectable collagen-Matrigel hydrogel mixtures were combined with CD56+ enriched myoblasts and injected into the PDMS-based chips to a final cell density of 15 and 20 million cells/ml, for young and old, and allowed to polymerize at 37°C for 60 min as previously described (Giza et al., 2022; Parafati et al., 2023).

Differentiation followed a two-step process as previously described (Giza et al., 2022; Parafati et al., 2023). Differentiation Media 1 included MEM-α, 0.5% (v/v) ITS, 2%(v/v) B27,10 μM DAPT, 1 μM Dabrafenib, 20 mM HEPES, pH 7.3 and, 0.1 mg/mL Primocin and Differentiation Media 2 included MEM-α, 0.5% (v/v) ITS, 2% (v/v) B-27, 20 mM HEPES, pH 7.3 and 0.1 mg/mL Primocin. Stage 2 medium was stored in a gas permeable bag for cold storage. Of the 16 chips, there were eight chip replicates of the YA mean cohort and eight chip replicates of the OS mean cohort (8xYA and 8x OS). Also, Space Tango’s CubeLab™ designed to maintain environmental controls beyond facility ambient conditions, captured analytics of the tissue chips. On day 14, thirty minutes of electrical stimulation (E-Stim) of 3 V, 2 Hz, 2msec pulse was applied sequentially to four YA and four OS tissue chips, the other eight chips (4xYA and 4x OS) were non-electrically stimulated (No E-Stim) (**Figure 1**). Donor-derived 3D myobundles were maintained in Space Tango’s autonomous CubeLab™ payload at 37°C and 5-6% CO_2_, and ground control samples were cultured and E-Stim in the same timeframe as flight with Space Tango’s CubeLab™. The myobundles were cultured in Stage II maintenance medium that was changed every 6 h to maintain healthy muscle microtissues, providing nutrients and gas exchange. Ground control samples were maintained identically, with media changes replicated exactly on a 6 h from the ISS.

### Measurement of Myotube Contractile Activity and Digital Image Correlation Analysis

Myotube contractility was quantified by digital image correlation (DIC), an approach that measures surface deformation. The DIC analysis was done by tracking the regional variation of gray-scale features between two succession images recorded before and after deformation, defined as static/undeformed and target/deformed images, respectively. Reference images were taken before and during E-Stim by an optics system installed in the CubeLab^TM^ as previously described (Parafati et al., 2023). Images are acquired with a resolution of 3088 × 2076 pixels (2.6 x 1.7 mm). A pixel corresponds to 0.84 µm. Image processing was similarly described with modifications (Giza et al., 2022). In brief, 40 sec videos captured at 40 fps in compressed video file formats (mp4) were extracted for a sequence of 1601 tiff images each using MATLAB software. DIC was performed using motion analyzer GOM correlate professional software (Zeiss, Germany) to determine the synchronicity of contractile oscillating pattern and displacement of the engineered myobundles’ region of interest (ROI) which included the entire image excluding the micro post). The reference image (the first frame prior to application of E-Stim image) is compared to each test image (i.e. with E-Stim applied) in the sequence. After the analysis, the DIC values are saved as a raw EXCEL file that stores the numerical displacement info for each grid at individual frames. Each frame represents a timepoint and subsequent frames are plotted as time on the X-axis. The average pixel value across the rows and columns in the ROI is converted to microns and plotted on the Y-axis as displacement. By this process, the 2D graph of displacement vs. time is generated.

### RNA isolation and preparation

We have performed transcript profiling of YA and OS myoblasts cultured for 2 days as well as of both YA and OS electrically stimulated (E-Stim) myotubes after 12 days of spaceflight and compared with ground-based analog studies by RNA-sequencing (RNA-seq). Myoblasts and myobundles cultured in tissue-on-chip devices were preserved with RNAlater® (ThermoFisher) and frozen down at −30°C in spaceflight and Earth (**Figure 1**). Once RNAlater® was removed, lysis buffer RLT Plus (Qiagen) plus β-mercaptoethanol was used for lysing myobundles in the chip. Then RNA isolation was performed using RNeasy Plus Mini Kit according to manufacturer’s instructions (Qiagen). RNA concentration and quality were determined with Agilent TapeStation 2200 (Agilent Technologies, Santa Clara, CA, USA) (**Supplementary table S1**). For each sample, 500 ng of total RNA were used as input material for RNA sample preparation. In total, there were 30 sequencing samples, each group had triplicate biological replicates. The sequencing of six RNA-seq libraries constructed from YA- and OS-derived myoblasts immediately before induction of myogenic differentiation and 24 RNA-seq libraries constructed from E-Stim and No E-Stim YA- and OS-derived myobundles on day 21 of induced myogenic differentiation generated more than 50 million unique mapped sequencing reads per library.

### RNA sequencing library preparation and RNA-Seq data processing

Strand-specific RNA-Seq libraries were generated using the NEBNext® Ultra™ Directional RNA Library Prep Kit for Illumina (NEB, USA) RNA sample preparation kit and yielded fragments with 220–700 base pairs. The qualified fragments were ligated with 60 adapters, amplified, and submitted for sequencing by Illumina NovaSeq 6000 (Illumina, San Diego, CA) to generate paired end reads with a length of 150 bases. The input sequences were trimmed using trimmomatic to remove Illumina adapters and reads shorter than 20 bp. Quality control was performed before and after trimming using FastQC (v 0.11.4) and MultiQC53 and a total of 50 million reads were generated for each sample yielding coverage in the range of 118 to 220 bases for each sample. Then the input sequences were aligned to the transcriptome using the STAR aligner, version 2.7.9a (54). Transcript concentration was quantified using RSEM (RSEM v1.3.1) (55), and genes with insufficient average counts (define number) were excluded from further statistical analysis. Differential expression analysis was performed using the DESeq2 package (56), with an FDR-corrected P-value threshold of 0.05. Sequencing and bioinformatics were performed at University of Florida Interdisciplinary Center for Biotechnology Research core facility (UF | ICBR).

### Functional enrichment analysis of DEGs

In this study, to facilitate the transition from data sets to biological meaning, RNA-seq data from young and old myobundles, exposed to microgravity and terrestrial conditions, were subjected to multiple approaches to calculate signaling pathway enrichments. To reveal functional enrichment of gene data sets we performed Gene Ontology (GO) analysis (Huang et al., 2009). Biological process (BP), molecular function (MF), and cellular component (CC) are the three categories under which GO enrichment was classified. The functional interpretation of genomic information was also related to Kyoto Encyclopedia of Genes and Genomes (KEGG) enrichment analysis (Kanehisa et al., 2000). GO annotation and KEGG pathway enrichment analysis were obtained utilizing bioinformatics tools from the curated Database for Annotation, Visualization, and Integrated Discovery (DAVID Gene Functional Classification Tool, https://david.ncifcrf.gov/). Furthermore, significant genes were grouped according to REACTOME databases (http://www.reactome.org), which annotates DEGs in pathway topological graphs to visualize involved genes, gene–gene interactions, and regulatory information in the pathway (Fabregat et al., 2017, 2018). To perform GO, KEGG and REACTOME analysis, we selected genes that were differentially expressed using FDR ≤ 0.05 and log_2_FC ≥ ±0.5 or ≥±1.5 cutoffs, as described in figure legends. For differential expression, thresholds of corrected enrichment p-value ≤ 1E-6 and p-value ≤ 0.01 were used for generating GO plots, as described in figure legends; and corrected enrichment p-value ≤ 0.05 was used for KEGG and REACTOME analysis. Only pathways with gene counts > 5 from the input list were significantly enriched. The Annotation Data Set was set to PANTHER Protein Class and analysis was performed using Fisher’s Exact Test with False Discovery Rate correction.

### Statistical analysis

Quality control, mapping, and expression for RNA-seq analysis were performed, and the Padj of genes was set to ≤ 0.05 to detect differentially expressed genes (DEGs). Data was analyzed and plotted using GraphPad Prism 8.4.3 (GraphPad Software Inc.), and all quantitative data are presented as the mean ± SD. In DAVID database, Fisher’s Exact test is adopted to measure the gene-enrichment in annotation terms and to identify the over-represented pathways with significant overlaps with pre-defined gene sets. Fisher’s Exact p-values are computed by summing probabilities p over defined sets of tables (Prob = ΣAp) and looks for any functional enrichment against a background that consist of all Homo Sapiens genes. Each GO term is used to describe the features of genes and gene products. For each Gene Ontology (GO) term, the number of DEGs annotated to the term is compared against the number of genes expected by chance. Significance of each GO term was assessed using the default homo sapiens GO annotation as background. A GO term was considered statistically significant at FDR-corrected p-value ≤ 1E-6 and p-value ≤ 0.01.

## Supporting information

YA ground video

YA flight video

OS flight video

OS ground video

## RESOURCE AVAILABILITY

### Lead Contact

Further information and requests for resources and reagents should be directed to the lead contact, Siobhan Malany.

### Materials Availability

This study did not generate new unique reagents.

Unless otherwise noted, data are available in the main text or the supplementary materials.

### Data and Code Availability

The accession number for the RNA-Sequencing data reported here is Gene Expression Omnibus: TBD. NASA GeneLab # TBD. Donor-derived cells are obtained through a materials transfer agreement and based on availability.

## ACKNOWLEDGMENTS

The authors acknowledge the team at Space Tango, Twyman Clements, Shelby Giza, Paul Gamble, Benjamin Lumpp, Isabel Moore, Jason Rexroat, and David Mays for assembly and performance of the CubeLab^TM^ that housed the experiment, The PAUL accent locker, and the on-orbit Tango Lab. We thank Don Platt of Micro Aerospace Solutions for assembling the electrode leads for the tissue chips. The Authors thank the scientific program manager Dr. Lucie Low for advocating the NIH Tissue Chips program at the National Center for Advancing Translational Sciences (NCATS). We thank payload operations project manager Erica Bumgardner for the research and development payloads utilizing the ISS U.S. National Laboratory. We thank Elizabeth Carter, Zach Lawson, and Anne Currin at Kennedy Space Center (KSC) - Space Station Processing Facility (SSPF) operations in Cape Canaveral. We thank the Interdisciplinary Center at the University of Florida for, Biotechnology Research core facility (UF | ICBR) for RNA sequencing. We thank Shayan Abbas for helping with digital image correlation analysis. We also acknowledge logistic support from Mauro Parlavecchio who helped the UF team at KSC laboratory and Mario Garcia from Micro-gRx, INC, for documenting our journey. We acknowledge the following funding sources: National Institutes of Health grant 5UG3TR002598 (S.M), National Institutes of Health grant 5UH3TR002598 (S.M.); National Institutes of Health grant 5UH3TR002598-05S1 Admin Suppl (S.M.); University of Florida Prosper Bridge Fund (M.P.); Center for Advancement of Science in Space #UA-2019-011 (S.M) and NASA task order to Space Tango (T.C.), Micro Aerospace Solutions (D.P.), and Micro-gRx (L.M).

## AUTHOR CONTRIBUTIONS

MP and SM conceived and designed the project. MP and SM performed and analyzed most experiments with assistance from ZT and PC. MP and SM performed experiments at Kennedy Space Center SSPF Labs with assistance from LM. PC led the clinical study from which donor derived myoblasts were derived. MP performed bioinformatics analysis and DIC analysis with assistance from ZT. SM supervised the project and acquired funding. SM and MP wrote the manuscript with input from all authors. SM, MP, and PC edited the manuscript.

