## Supplemental material for "Microgravity Accelerates Skeletal Muscle Degeneration: Functional and Transcriptomic Insights from a Muscle Lab-on-Chip Model Onboard the ISS"

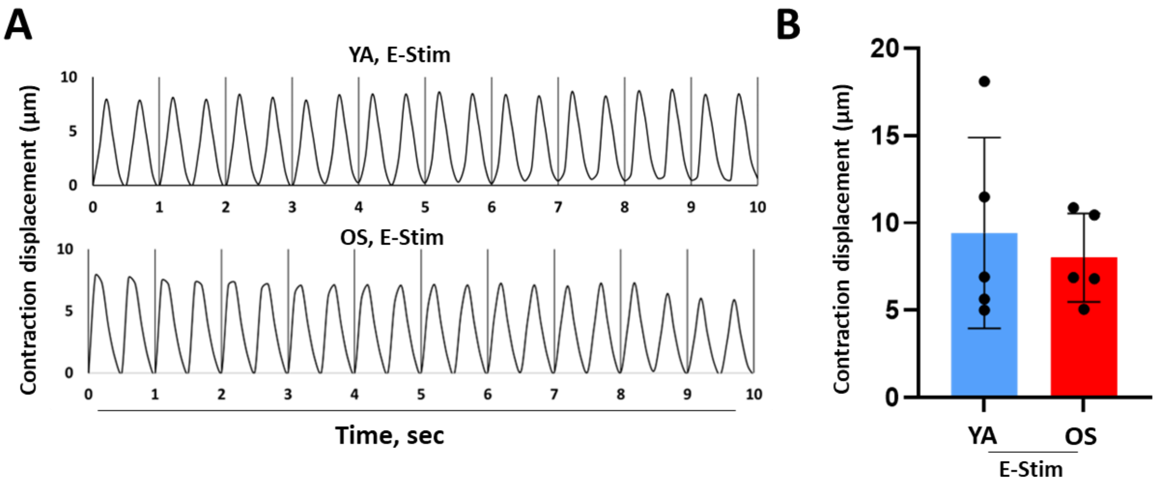


**Figure S1. Myobundles from YA- and OS-donor derived cells exhibit similar contractile function.** (A) A 10 sec timeframe of contraction displacement responses for five OS- and five YA-derived myobundles under 3 V, 2 Hz, 2 ms electric field as determined by digital image correlation analysis. Vidoes were recrded for 30 min at 10 fps (B) Bar graph of average peak heights of contraction displacement responses determined over 40 sec timeframes for OS- (red) and YA-(blue) derived myobundles (N= average of 5 myobundles per group). As the reference contraction displacement, donor-derived myobundles were recorded for 40 sec without E-Stim. The contraction displacement values were zero.

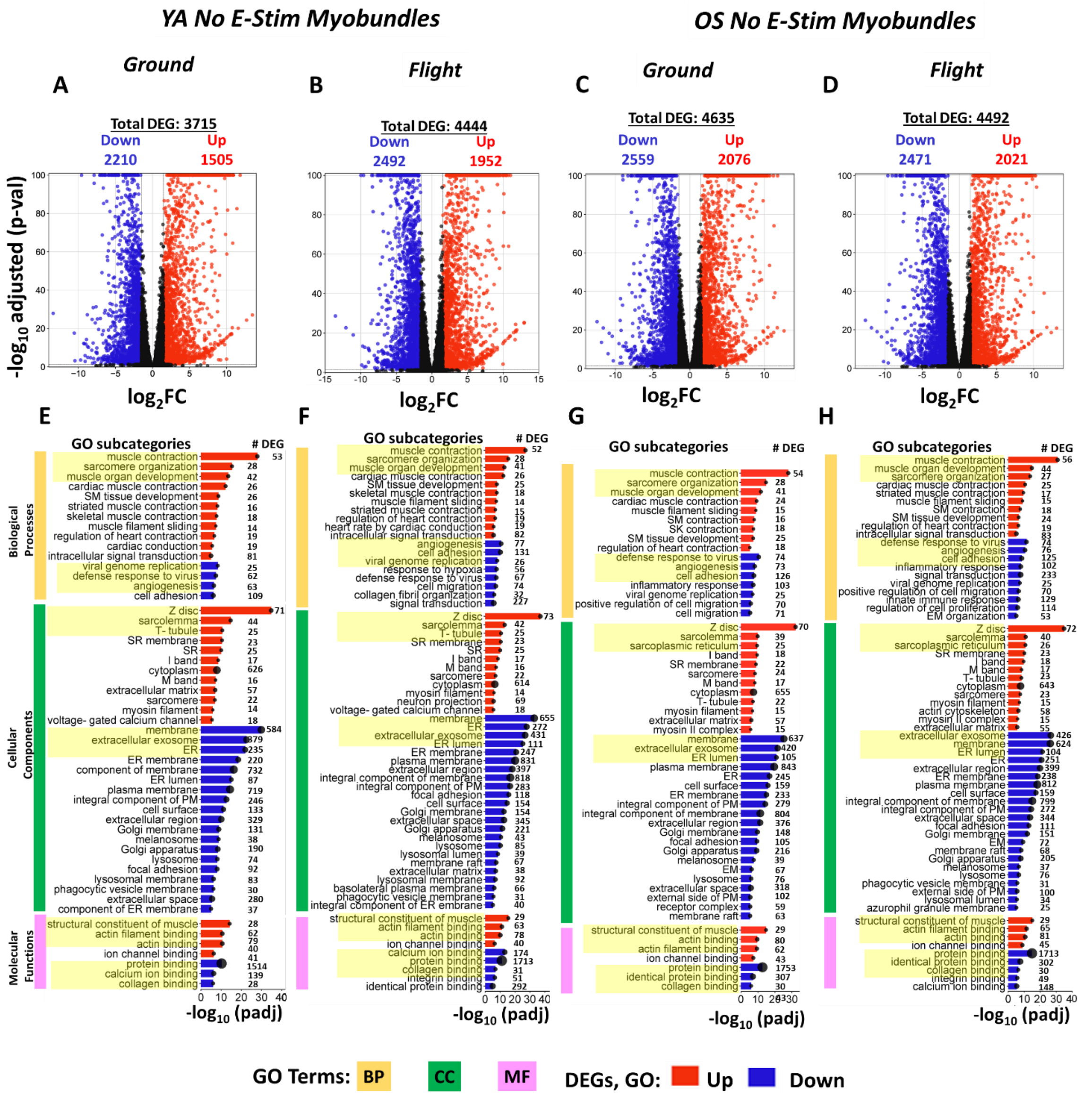

**Figure S2. RNA-Seq analysis of DEGs comparing flight and ground No E-Stim on day 21 vs. day 2**. Global gene expression changes of log2 fold change ($log_2FC$) versus –log10 fold discovery rate (FDR) for **(A)** YA ground, **(B)** YA flight, **(C)** OS ground, and **(D)** OS flight. Red and blue circles indicated increased and decreased activity of DEGs, respectively, with at least FDR ≤ 0.05 and $log_2FC \geq \pm 1.5$ cutoff. Black circles denote DEGs below the cutoff. Gene ontology (GO) enrichment analysis is shown for **(E)** YA ground, **(F)** YA flight, **(G)** OS ground, and **(H)** OS flight for biological processes (BP), cellular component (CC) and molecular function (MF), ranked by log10 FDR adjusted p-values, was performed using DAVID gene functional classification tool on the DEGs (FDR≤0.05 and $log_2FC \geq\pm 1.5$) between the indicated groups based on a two-tailed Student's t test. The horizontal axis represents the log10 adjusted p values of DEGs, while the vertical axis represents enriched subcategories. Count (circle size) indicates the number of enriched DEGs reported per subcategory. Bonferroni corrected p-value for each GO subcategory was set as threshold ≤ 0.01. Data are representative of three independent tissue chip RNA-seq determinations. Yellow color highlights the top three gene categories with the highest adjusted p-value (padj).

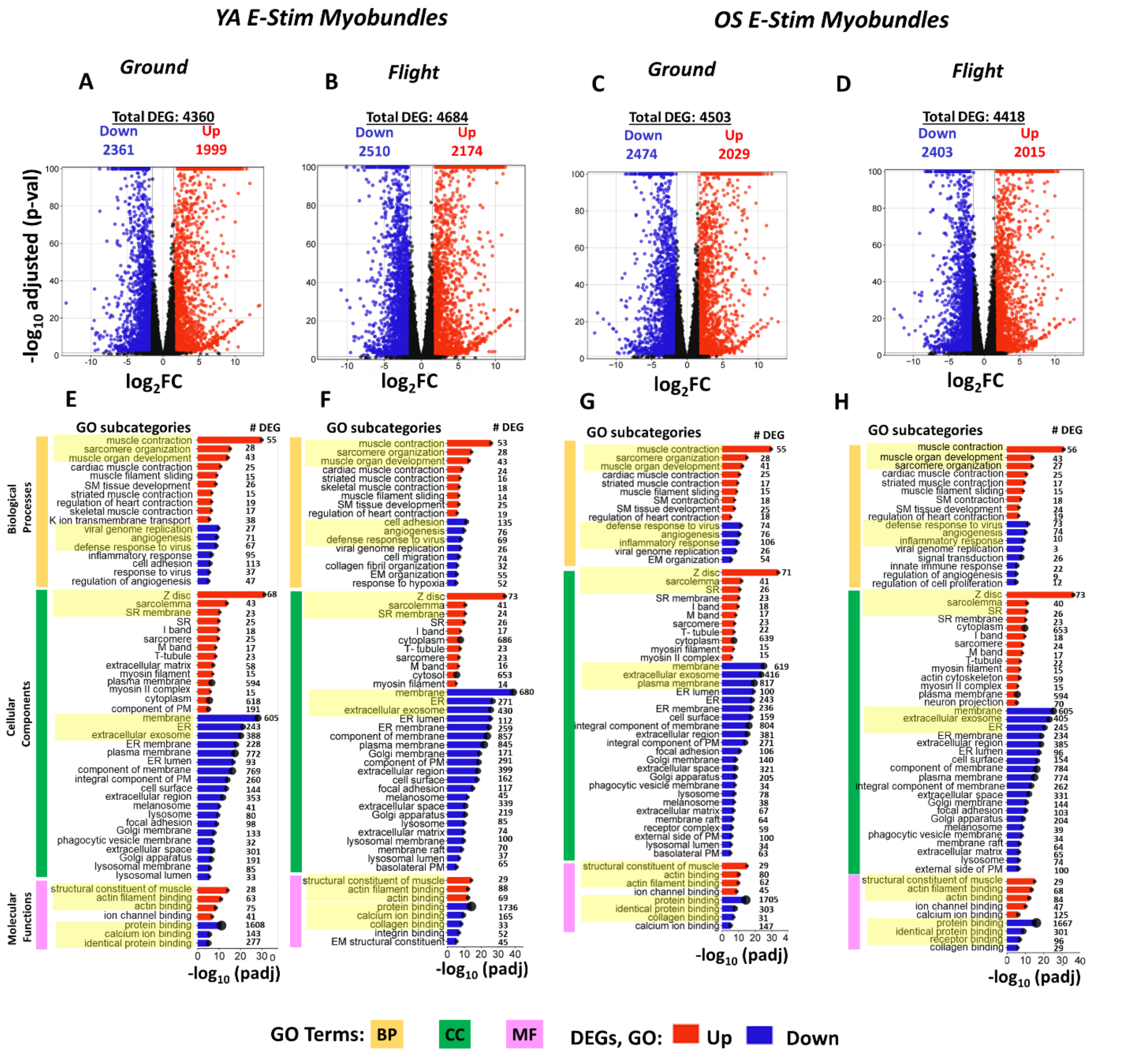

**Figure S3. RNA-Seq analysis of DEGs comparing flight and ground E-Stim tissue chips on day 21 vs. day 2**. Global gene expression changes of log2 fold change (log2FC) versus –log10 fold discovery rate (FDR) for **(A)** YA ground, **(B)** YA flight, **(C)** OS ground, and **(D)** OS flight. Red and blue circles indicated increased and decreased activity of DEGs, respectively, with at FDR ≤ 0.05 and log2FC ≥ ± 1.5 cutoff. Black circles denote DEGs below the cutoff. Gene ontology (GO) enrichment analysis is shown for **(E)** YA ground, **(F)** YA flight, **(G)** OS ground, and **(H)** OS flight for biological processes (BP), cellular component (CC) and molecular function (MF), ranked by log10 FDR adjusted p-values, was performed using DAVID gene functional classification tool (FDR ≤ 0.05 and $\log_2$FC ≥± 1.5) between the indicated groups based on a two-tailed Student's t test. The horizontal axis represents the log10 adjusted p values of enriched DEGs to background genes, while the vertical axis represents enriched subcategories. Count (circles) indicates the number of DEGs reported in each subcategory. Bonferroni corrected p-value was set as threshold ≤ 0.01. Data are representative of three independent tissue chip RNA-seq determinations. Yellow color highlights the top 3 increased or decreased gene categories with the highest adjusted p-value (padj).

**Table S1. RNA quality and quantity for myobundles in spaceflight and ground**

| Groups | Sample # | Chip # | Description | RNA (ng) | RIN |
|---|---|---|---|---|---|
| **Flight differentiated myobundles** | | | | | |
| 1 | 1 | 1 | Young E-Stim | 3.906 | 9.6 |
| | 2 | 2 | Young E-Stim | 3.192 | 8.8 |
| | 3 | 3 | Young E-Stim | 4.5804 | 9.1 |
| 2 | 4 | 5 | Old E-Stim | 5.313 | 9.7 |
| | 5 | 6 | Old E-Stim | 6.209 | 9.7 |
| | 6 | 7 | Old E-Stim | 4.8246 | 9.4 |
| 3 | 7 | 9 | Young No E-Stim | 3.5838 | 9.4 |
| | 8 | 10 | Young No E-Stim | 4.4712 | 9.1 |
| | 9 | 11 | Young No E-Stim | 3.8657 | 9.7 |
| 4 | 10 | 13 | Old No E-Stim | 4.3007 | 9.6 |
| | 11 | 14 | Old No E-Stim | 5.355 | 9.7 |
| | 12 | 15 | Old No E-Stim | 5.8106 | 9.6 |
| **Ground differentiated myobundles** | | | | | |
| 5 | 13 | 2 | Young E-Stim | 3636.81 | 9.5 |
| | 14 | 3 | Young E-Stim | 3249 | 9.1 |
| | 15 | 4 | Young E-Stim | 3244.17 | 9.4 |
| 6 | 16 | 6 | Old E-Stim | 3841.11 | 9.6 |
| | 17 | 7 | Old E-Stim | 3225.09 | 9.3 |
| | 18 | 8 | Old E-Stim | 3404.76 | 9.3 |
| 7 | 19 | 9 | Young No E-Stim | 3180.93 | 9 |
| | 20 | 11 | Young No E-Stim | 3718.2 | 8.8 |
| | 21 | 12 | Young No E-Stim | 3202.53 | 9.3 |
| 8 | 22 | 13 | Old No E-Stim | 4209.33 | 9.2 |
| | 23 | 15 | Old No E-Stim | 2767.62 | 9.2 |
| | 24 | 16 | Old No E-Stim | 3021.84 | 9.5 |
| **Undifferentiated myobundles** | | | | | |
| 9 | 25 | 1 | Young day 2 | 2786.941 | 9.6 |
| | 26 | 2 | Young day 2 | 2690.881 | 9.5 |
| | 27 | 3 | Young day 2 | 2684.554 | 9.6 |
| 10 | 28 | 4 | Old day 2 | 2687.754 | 9.7 |
| | 29 | 5 | Old day 2 | 3173.305 | 9.6 |
| | 30 | 6 | Old day 2 | 3174.541 | 9.6 |

**Table S2. RNA-seq. data analysis of muscle age-related genes** visualized as $log_2$FC mean values for the E-Stim young vs. old-derived myobundles comparisons in gravity and microgravity (Su et al., 2015). Data are representative of three independent RNA-seq. determinations.

| **Table 1** | | | | | |
|---|---|---|---|---|---|
| **Gene** | **Gene name** | ***YA No ES vs. OS No ES*** | | ***YA ES vs. OS ES*** | |
| | **Upregulated aging genes** | **Gravity** | **Microgravity** | **Gravity** | **Microgravity** |
| **ALOX5AP** | **arachidonate 5-lipoxygenase activating protein** | **0.00** | **0.00** | **0.00** | **2.50** |
| **CCR10** | **C-C motif chemokine receptor 10** | **0.00** | **0.00** | **0.00** | **1.90** |
| **SAT1** | **spermidine/spermine N1-acetyltransferase 1** | **0.00** | **0.00** | **0.00** | **1.72** |
| **CFD** | **complement factor D** | **0.00** | **0.00** | **0.00** | **1.30** |
| **DYNC1H1** | **dynein cytoplasmic 1 heavy chain 1** | **0.00** | **0.00** | **0.00** | **1.27** |
| **IFRD1** | **interferon related developmental regulator 1** | **0.00** | **0.00** | **0.00** | **1.21** |
| **ASS1** | **argininosuccinate synthase 1** | **0.00** | **0.00** | **0.00** | **1.19** |
| **WISP2** | **WNT1 inducible signaling pathway 2** | **0.00** | **0.00** | **0.00** | **1.19** |
| **ABRA** | **actin binding Rho activating protein** | **0.00** | **0.00** | **0.00** | **1.11** |
| **CLK4** | **CDC like kinase 4** | **0.00** | **-0.81** | **0.71** | **1.06** |
| **FGGY** | **FGGY carbohydrate kinase domain containing** | **0.00** | **0.00** | **0.00** | **1.05** |
| **NENF** | **neudesin neurotrophic factor** | **0.00** | **0.00** | **0.00** | **1.05** |
| **AKR1C3** | **aldo-keto reductase family 1 member C3** | **0.00** | **0.00** | **0.00** | **0.98** |
| **TRMT112** | **tRNA methyltransferase activator subunit 11-2** | **0.00** | **0.00** | **0.00** | **0.91** |
| **TPPP3** | **tubulin polymerization promoting protein family member 3** | **0.00** | **0.00** | **0.00** | **0.90** |
| **RPS27L** | **ribosomal protein S27 like** | **0.00** | **0.00** | **0.00** | **0.90** |
| **IRS2** | **insulin receptor substrate 2** | **0.00** | **0.00** | **0.00** | **0.88** |
| **AHNAK** | **AHNAK nucleoprotein** | **0.00** | **0.00** | **0.00** | **0.87** |
| **BTF3** | **basic transcription factor 3** | **0.00** | **0.00** | **0.00** | **0.84** |
| **SFPQ** | **splicing factor proline and glutamine rich** | **0.81** | **0.00** | **0.00** | **0.84** |
| **MMD** | **monocyte to macrophage differentiation associated** | **0.00** | **0.96** | **0.00** | **0.79** |
| **ECHDC2** | **enoyl-CoA hydratase domain containing 2** | **0.00** | **0.00** | **0.00** | **0.79** |
| **DUSP1** | **dual specificity phosphatase 1** | **0.93** | **0.00** | **0.00** | **0.78** |
| **KARS** | **Lysine--tRNA ligase** | **0.00** | **0.00** | **0.00** | **0.77** |
| **DDB2** | **damage specific DNA binding protein 2** | **0.00** | **0.00** | **0.00** | **0.75** |
| **SAP30BP** | **SAP30 binding protein** | **0.00** | **0.00** | **0.00** | **0.74** |
| **CCDC28A** | **coiled-coil domain containing 28A** | **0.00** | **0.00** | **0.00** | **0.73** |

| | | | | | |
|---|---|---|---|---|---|
| **MYL6B** | **myosin light chain 6B** | **0.00** | **0.00** | **0.00** | **0.72** |
| **PXDC1** | **PX domain containing 1** | **0.00** | **0.00** | **0.67** | **0.70** |
| **RPL22** | **ribosomal protein L22** | **0.00** | **0.00** | **0.00** | **0.70** |
| **NPRL2** | **NPR2 like, GATOR1 complex subunit** | **0.00** | **0.00** | **0.00** | **0.69** |
| **DNAJC8** | **DnaJ heat shock protein family (Hsp40) member C8** | **0.00** | **0.00** | **0.00** | **0.68** |
| **ATXN1** | **ataxin 1** | **0.00** | **0.00** | **0.00** | **0.67** |
| **RFXANK** | **regulatory factor X associated ankyrin containing protein** | **0.00** | **0.00** | **0.00** | **0.64** |
| **SIRT1** | **sirtuin 1** | **0.00** | **0.00** | **0.00** | **0.62** |
| **GBP2** | **guanylate binding protein 2** | **0.00** | **0.00** | **0.00** | **0.60** |
| **LUC7L3** | **LUC7 like 3 pre-mRNA splicing factor** | **0.00** | **0.00** | **0.00** | **0.58** |
| **CAMLG** | **calcium modulating ligand** | **0.00** | **0.00** | **0.00** | **0.58** |
| **KLF4** | **KLF transcription factor 4** | **0.00** | **0.00** | **0.00** | **0.58** |
| **NOSIP** | **nitric oxide synthase interacting protein** | **0.00** | **0.00** | **0.00** | **0.56** |
| **NAP1L1P3** | **nucleosome assembly protein 1 like 1 pseudogene 3** | **0.00** | **0.00** | **0.00** | **0.56** |
| **VPS13A** | **vacuolar protein sorting 13 homolog A** | **0.00** | **0.00** | **0.00** | **0.56** |
| **CLTA** | **clathrin light chain A** | **0.00** | **0.00** | **0.00** | **0.55** |
| **ILKAP** | **ILK associated serine/threonine phosphatase** | **0.00** | **0.00** | **0.00** | **0.53** |
| **POP7** | **POP7 homolog, ribonuclease P/MRP subunit** | **0.00** | **0.00** | **0.00** | **0.51** |
| **ABCC5** | **ATP binding cassette subfamily C member 5** | **0.68** | **0.00** | **0.00** | **0.00** |
| **ADAMTS1** | **ADAM metallopeptidase with thrombospondin type 1 motif 1** | **0.00** | **-0.66** | **0.00** | **0.00** |
| **ADI1** | **acireductone dioxygenase 1** | **-0.61** | **0.00** | **-0.57** | **0.00** |
| **CACNG1** | **calcium voltage-gated channel auxiliary subunit gamma 1** | **-0.63** | **0.00** | **-0.62** | **0.00** |
| **CLTCL1** | **clathrin heavy chain like 1** | **-0.71** | **0.00** | **0.00** | **0.00** |
| **CTNNA2** | **catenin alpha 2** | **0.00** | **0.00** | **1.87** | **0.00** |
| **DCN** | **decorin** | **0.87** | **0.00** | **0.00** | **0.00** |
| **DDX3Y** | **DEAD-box helicase 3 Y-linked** | **0.78** | **0.00** | **0.81** | **0.00** |
| **DECR2** | **2,4-dienoyl-CoA reductase 2** | **-0.89** | **-0.84** | **-0.84** | **0.00** |
| **EPB41L3** | **erythrocyte membrane protein band 4.1 like 3** | **0.00** | **0.00** | **-0.84** | **0.00** |
| **FANCE** | **FA complementation group E** | **-0.81** | **0.00** | **-0.57** | **0.00** |
| **GDI1** | **GDP dissociation inhibitor 1** | **0.69** | **0.00** | **0.00** | **0.00** |
| **HIST1H2BK** | **H2B clustered histone 12** | **-0.52** | **0.00** | **-0.54** | **0.00** |
| **HNRNPH3** | **heterogeneous nuclear ribonucleoprotein H3** | **0.57** | **0.00** | **0.00** | **0.00** |
| **IL15** | **interleukin 15** | **-1.13** | **0.00** | **-0.88** | **0.00** |

| | | | | | |
|---|---|---|---|---|---|
| **KLF10** | **KLF transcription factor 10** | **1.08** | **0.00** | **0.99** | **0.00** |
| **KLF11** | **KLF transcription factor 11** | **0.00** | **0.00** | **0.55** | **0.00** |
| **LGI1** | **leucine rich glioma inactivated 1** | **-0.70** | **0.00** | **0.00** | **0.00** |
| **LTBP2** | **latent transforming growth factor beta binding protein 2** | **1.16** | **0.00** | **1.42** | **0.00** |
| **MT3** | **metallothionein 3** | **-1.45** | **0.00** | **0.00** | **0.00** |
| **MYOM2** | **myomesin 2** | **-0.90** | **0.00** | **0.00** | **0.00** |
| **PFKFB2** | **6-phosphofructo-2-kinase/fructose-2,6-biphosphatase 2** | **0.00** | **0.00** | **0.69** | **0.00** |
| **PIK3R1** | **phosphoinositide-3-kinase regulatory subunit 1** | **0.81** | **0.00** | **0.58** | **0.00** |
| **PLA1A** | **phospholipase A1 member A** | **1.08** | **0.00** | **0.00** | **0.00** |
| **POMT1** | **protein O-mannosyltransferase 1** | **0.00** | **0.00** | **0.00** | **0.00** |
| **RBM28** | **RNA binding motif protein 28** | **0.00** | **0.00** | **0.53** | **0.00** |
| **SLC7A2** | **solute carrier family 7 member 2** | **-0.64** | **-0.88** | **0.00** | **0.00** |
| **SLC7A6** | **solute carrier family 7 member 6** | **0.00** | **0.00** | **0.87** | **0.00** |
| **SYNPO** | **synaptopodin** | **-0.52** | **0.00** | **0.00** | **0.00** |
| **SYNRG** | **synergin gamma** | **0.93** | **0.00** | **0.00** | **0.00** |
| **ZNF136** | **zinc finger protein 136** | **0.00** | **1.29** | **0.00** | **0.00** |
| **MTHFSD** | **methenyltetrahydrofolate synthetase domain containing** | **0.00** | **0.00** | **0.00** | **-0.51** |
| **CDC42EP3** | **CDC42 effector protein 3** | **0.00** | **0.00** | **0.00** | **-0.51** |
| **CELF2** | **CUGBP Elav-like family member 2** | **0.00** | **0.00** | **0.00** | **-0.52** |
| **XPC** | **XPC complex subunit, DNA damage recognition and repair factor** | **0.00** | **0.00** | **0.00** | **-0.52** |
| **BLCAP** | **BLCAP apoptosis inducing factor** | **-0.60** | **0.00** | **-0.52** | **-0.53** |
| **ARFGAP2** | **ADP ribosylation factor GTPase activating protein 2** | **0.00** | **0.00** | **0.00** | **-0.53** |
| **DDX5** | **DEAD-box helicase 5** | **0.51** | **0.00** | **0.00** | **-0.54** |
| **NR2C2** | **nuclear receptor subfamily 2 group C member 2** | **0.00** | **0.00** | **0.50** | **-0.54** |
| **TNS3** | **tensin 3** | **0.00** | **0.00** | **0.00** | **-0.56** |
| **PTBP1** | **polypyrimidine tract binding protein 1** | **0.00** | **0.00** | **0.00** | **-0.56** |
| **KAT6A** | **lysine acetyltransferase 6A** | **0.00** | **0.00** | **0.00** | **-0.59** |
| **GLG1** | **golgi glycoprotein 1** | **0.00** | **0.00** | **0.00** | **-0.61** |
| **MTSS1** | **MTSS I-BAR domain containing 1** | **0.00** | **0.00** | **0.00** | **-0.61** |
| **SF1** | **splicing factor 1** | **0.00** | **0.00** | **0.00** | **-0.62** |
| **MAN2B2** | **mannosidase alpha class 2B member 2** | **0.00** | **0.00** | **0.00** | **-0.65** |
| **LAMP1** | **lysosomal associated membrane protein 1** | **0.00** | **0.00** | **0.00** | **-0.67** |
| **TRIM8** | **tripartite motif containing 8** | **0.00** | **0.00** | **0.00** | **-0.68** |
| **WARS** | **tryptophanyl-tRNA synthetase** | **0.00** | **0.00** | **0.00** | **-0.70** |
| **FOXO1** | **forkhead box O1** | **0.00** | **0.00** | **0.00** | **-0.70** |

| | | | | | |
|---|---|---|---|---|---|
| **MAGED1** | **MAGE family member D1** | **-0.59** | **0.00** | **-0.65** | **-0.71** |
| **SLIT2** | **slit guidance ligand 2** | **0.00** | **0.00** | **0.00** | **-0.73** |
| **SIDT2** | **SID1 transmembrane family member 2** | **0.00** | **0.00** | **0.00** | **-0.74** |
| **METTL7A** | **methyltransferase like 7A** | **-0.88** | **-0.72** | **-0.96** | **-0.75** |
| **LRRC8D** | **leucine rich repeat containing 8 VRAC subunit D** | **0.00** | **0.00** | **0.00** | **-0.76** |
| **ZNF134** | **zinc finger protein 134** | **0.00** | **0.00** | **0.00** | **-0.78** |
| **ADAMTS5** | **ADAM metallopeptidase with thrombospondin type 1 motif 5** | **0.00** | **0.00** | **0.00** | **-0.82** |
| **PSMD5** | **proteasome 26S subunit, non-ATPase 5** | **0.00** | **0.00** | **0.00** | **-0.85** |
| **H6PD** | **hexose-6-phosphate dehydrogenase/glucose 1-dehydrogenase** | **0.00** | **0.00** | **0.00** | **-0.85** |
| **SPOCK1** | **SPARC (osteonectin), cwcv and kazal like domains proteoglycan 1** | **0.00** | **0.00** | **0.74** | **-0.90** |
| **SERPING1** | **serpin family G member 1** | **0.00** | **0.00** | **0.00** | **-0.91** |
| **LMO3** | **LIM domain only 3** | **-0.91** | **-0.84** | **0.00** | **-0.92** |
| **AKAP12** | **A-kinase anchoring protein 12** | **1.00** | **0.00** | **0.00** | **-0.92** |
| **TECPR2** | **tectonin beta-propeller repeat containing 2** | **0.00** | **0.00** | **0.00** | **-0.95** |
| **ANTXR2** | **ANTXR cell adhesion molecule 2** | **0.00** | **0.00** | **0.00** | **-0.96** |
| **LRP1B** | **LDL receptor related protein 1B** | **-1.26** | **0.00** | **-1.20** | **-1.03** |
| **ADAR** | **adenosine deaminase RNA specific** | **0.63** | **0.00** | **0.00** | **-1.03** |
| **TGFBR3** | **transforming growth factor beta receptor 3** | **1.04** | **0.00** | **1.04** | **-1.13** |
| **GLIS3** | **GLIS family zinc finger 3** | **0.00** | **0.00** | **1.41** | **-1.16** |
| **CACNA1E** | **calcium voltage-gated channel subunit alpha1 E** | **0.00** | **0.00** | **0.00** | **-1.41** |
| **TLR4** | **toll like receptor 4** | **0.00** | **0.00** | **0.00** | **-2.08** |
| **ABCA8** | **ATP binding cassette subfamily A member 8** | **0.00** | **-1.94** | **-1.49** | **-2.24** |
